# Kinase-modulated bioluminescent indicators enable noninvasive imaging of drug activity in the brain

**DOI:** 10.1101/2022.12.30.522361

**Authors:** Yan Wu, Joel R. Walker, Michael Westberg, Lin Ning, Michelle Monje, Thomas A. Kirkland, Michael Z. Lin, Yichi Su

## Abstract

Aberrant kinase activity contributes to the pathogenesis of brain cancers, neurodegeneration, and neuropsychiatric diseases, but identifying kinase inhibitors that function in the brain is challenging. Drug levels in blood do not predict efficacy in the brain because the blood-brain barrier prevents entry of most compounds. Rather, assessing kinase inhibition in the brain requires tissue dissection and biochemical analysis, a time-consuming and resource-intensive process. Here, we report kinase-modulated bioluminescent indicators (KiMBIs) for non-invasive longitudinal imaging of drug activity in the brain based on a recently optimized luciferase-luciferin system. We develop an ERK KiMBI to report inhibitors of the Ras-Raf-MEK-ERK pathway, for which no bioluminescent indicators previously existed. ERK KiMBI discriminates between brain-penetrant and non-penetrant MEK inhibitors, reveals blood-tumor barrier leakiness in xenograft models, and reports MEK inhibitor pharmacodynamics in native brain tissues and intracranial xenografts. Finally, we use ERK KiMBI to screen ERK inhibitors for brain efficacy, identifying temuterkib as a promising brain-active ERK inhibitor, a result not predicted from chemical characteristics alone. Thus, KiMBIs enable the rapid identification and pharmacodynamic characterization of kinase inhibitors suitable for treating brain diseases.

## Introduction

Aberrant kinase activity drives pathogenesis of multiple diseases of the central nervous system, including primary brain neoplasms and metastatic cancers in the brain^1–4^, neurodegenerative disorders such as Alzheimer’s and Parkinson’s diseases^5, 6^, and psychiatric disorders such as bipolar disease and schizophrenia^7^. However, most kinase pathway inhibitors previously developed to treat disorders outside the brain do not efficiently cross the blood-brain barrier (BBB)^1, 3, 6^. As a result, there is intense interest in developing new drugs to effectively inhibit specific kinase pathways in the brain^1, 3, 6^.

In typical drug discovery efforts, candidate molecules with appropriate in vitro potency and selectivity are screened for suitable pharmacokinetics prior to commencing expensive in vivo efficacy studies. For indications outside of the brain, pharmacokinetics is typically assessed by measuring drug concentrations in blood at different times after administration. However, drug concentrations in blood differ from concentrations in the brain due to the BBB^8^. Thus, in the case of brain targets, pharmacokinetics requires obtaining brain tissue at various times after drug administration, which is a terminal low-throughput procedure. An alternative way to determine drug concentrations in the brain would be to label candidate drugs with a radioactive element for imaging by PET or SPECT^9^, but this requires a synthetic pathway to incorporate the isotope, plus specialized expertise and equipment.

Another fundamental challenge is that pharmacokinetics can differ from pharmacodynamics, i.e., how the drug target responds over time. Cell type-specific drug export or metabolism may limit drug efficacy even with high drug levels in the surrounding tissue^10^. Accurate assessment of kinase inhibition can be especially problematic for brain cancers, where cancer cells can be dispersed amongst a larger number of normal cells. One solution is to perform immunocytochemistry with phosphorylation-specific antibodies to specific kinase substrates and with cancer-specific markers, allowing some determination of phosphorylation levels specifically in cancer cells^11^. However, this approach is highly time-consuming, resource-intensive, and requires high-quality phosphorylation-specific antibodies that may not exist for every drug target. Given these limitations of current methods to assess pharmacodynamics in the brain, a fast and inexpensive approach to evaluate kinase inhibitor activity within target cells in the brain would be highly desirable^12^.

Noninvasive imaging of kinase activity with genetically encoded biosensors could be one way to assess kinase inhibitor pharmacodynamics within target tissues in small animal models. To provide a near-real-time report of target engagement, the indicator kinetics with similar timescales as drug pharmacokinetics are needed, making transcription-based reporters unsuitable. Fluorescent protein-based reporters of kinase activity respond rapidly^13^, but fluorescence is poorly suited for non-invasive quantitative imaging through tissue. A more suitable approach is to engineer a bioluminescent enzyme whose activity can be directly modulated by kinase activity, as bioluminescence can be detected from essentially any location in the mouse brain with high signal-to-background ratios^14^.

However, past bioluminescent reporters of kinase activity were based on firefly luciferase (FLuc)^15, 16^, which introduces three limitations. First, FLuc produces an order of magnitude less light in vivo compared to modern luciferases^17^. Second, FLuc catalysis requires ATP, and photon emission from FLuc in animals has been shown to vary with conditions that alter ATP levels^18^. Thus a FLuc-based kinase reporter could respond artifactually to an off-target effect of a test compound on ATP levels via either a different kinase or a nonkinase pathway. Even on-target effects on kinases could produce artifactual responses from FLuc-based reporters, as suppression of kinase activity can induce either feedback activation of glucose metabolism over hours^19^ or reduced metabolism over days^20^. Thus, a FLuc-based kinase reporter may respond to both longterm effects of kinase inhibition on ATP, in addition to responding to kinase inhibition directly. Finally, FLuc appears especially prone to catalytic inhibition or protein stabilization by a wide variety of drugs. For example, 22 of 367 ATP-competitive kinase inhibitors were found to inhibit FLuc at sub-micromolar concentrations, whereas none inhibited the ATP-independent *Renilla* luciferase at these concentrations^21^. Different classes of drugs bind to FLuc outside the ATP-binding pocket to stabilize the protein, leading to increased bioluminescent signals over time in cells expressing FLuc^22^. The possibility of FLuc inhibition or stabilization by drug candidates is highly undesirable for measuring pharmacodynamics at extended timepoints.

Besides developing FLuc-independent bioluminescent indicators that are less prone to artifacts, indicators for inhibitors of the Ras-Raf-MEK-ERK pathway would be especially useful. This pathway (hereafter the Ras-ERK pathway) is hyperactivated in a large fraction of solid tumors, many of which metastasize to the brain^3, 4^. For instance, mutations are commonly observed in the upstream receptor tyrosine kinases EGFR and HER2 in brain metastases of lung and breast cancer, and in B-Raf in melanoma^4^. No inhibitors of Ras-ERK pathway components have been specifically approved for brain tumors, although numerous ones are under clinical investigation^1–4, 23^. Inhibitors approved for extracranial cancers can be administered to patients with brain metastases of the primary tumors, but results are typically mixed, and only recently have clinical trials specifically for brain metastases been performed^4^. Thus, rapid assessment of the intracranial activity of Ras-ERK pathway inhibitors in vivo would enable identification of compounds with the highest likelihood of clinical efficacy. However, surprisingly, no bioluminescent indicator of the Ras-ERK pathway of any type has been reported in the literature, whether based on FLuc or other luciferases.

Here, we report the development of kinase-modulated bioluminescent indicators (KiMBIs) for real-time visualization of kinase inhibition in living animals, including in the brain, through molecular engineering of the ATP-independent luciferase NanoLuc. Importantly, a substrate with improved performance in the brain was recently developed for NanoLuc, generating brain signals an order of magnitude brighter than achievable with FLuc (Su et al., in press). Given the importance of the Ras-ERK pathway in brain malignancies, we focused on developing a KiMBI for ERK, achieving a reporter that brightens 10-fold in response to ERK pathway inhibition. ERK KiMBI expressed in the mouse brain differentiated between brain permeant and impermeant inhibitors of the ERK activator mitogen-activated protein kinase kinase (MAPKK or MEK). ERK KiMBI also revealed that the BBB is disrupted in common xenograft models of glioma. In addition, KiMBI could track the pharmacodynamics of kinase inhibition over time in the same mouse, whereas pharmacodynamics using traditional tissue sampling methods would require a large number of animals. Finally, we used KiMBI to characterize the in-brain efficacy of ERK inhibitors in clinical trials. We identify LY3214996 (temuterkib) as a brain-active ERK inhibitor and discover that theoretically predicted BBB permeability and empirically determined in-brain efficacy are uncorrelated.

## Results

### Design and development of KiMBIs

To develop reporters that produce light upon kinase inhibition, we hypothesized that the catalytic activity of NanoLuc luciferase could be modulated through interactions between phosphorylated substrates (pSub) and phosphopeptide-binding domains (PBDs). The NanoBiT version of NanoLuc can be split after amino acid 156 in the loop before the last beta-strand, producing complementing fragments LgBiT and SmBiT, with SmBiT available as different affinity variants^24^. We postulated that inserting a PBD (or kinase substrate) between LgBiT and SmBiT, and fusing the cognate substrate (or PBD) elsewhere in the protein, could confer dephosphorylation-dependence on the enzymatic activity of NanoBiT. Specifically, when a phosphorylating kinase is highly active, the PBD-pSub interaction could tether the SmBiT in a conformation incompatible with binding to LgBiT, effectively competing with the weaker LgBiT-SmBiT interaction. In contrast, if the kinase or an upstream kinase is inhibited by a kinase inhibitor, then ongoing dephosphorylation could shift the equilibrium of conformational states toward LgBiT-SmBiT interaction, resulting in light production (**Fig. 1a**).

**Fig. 1.**
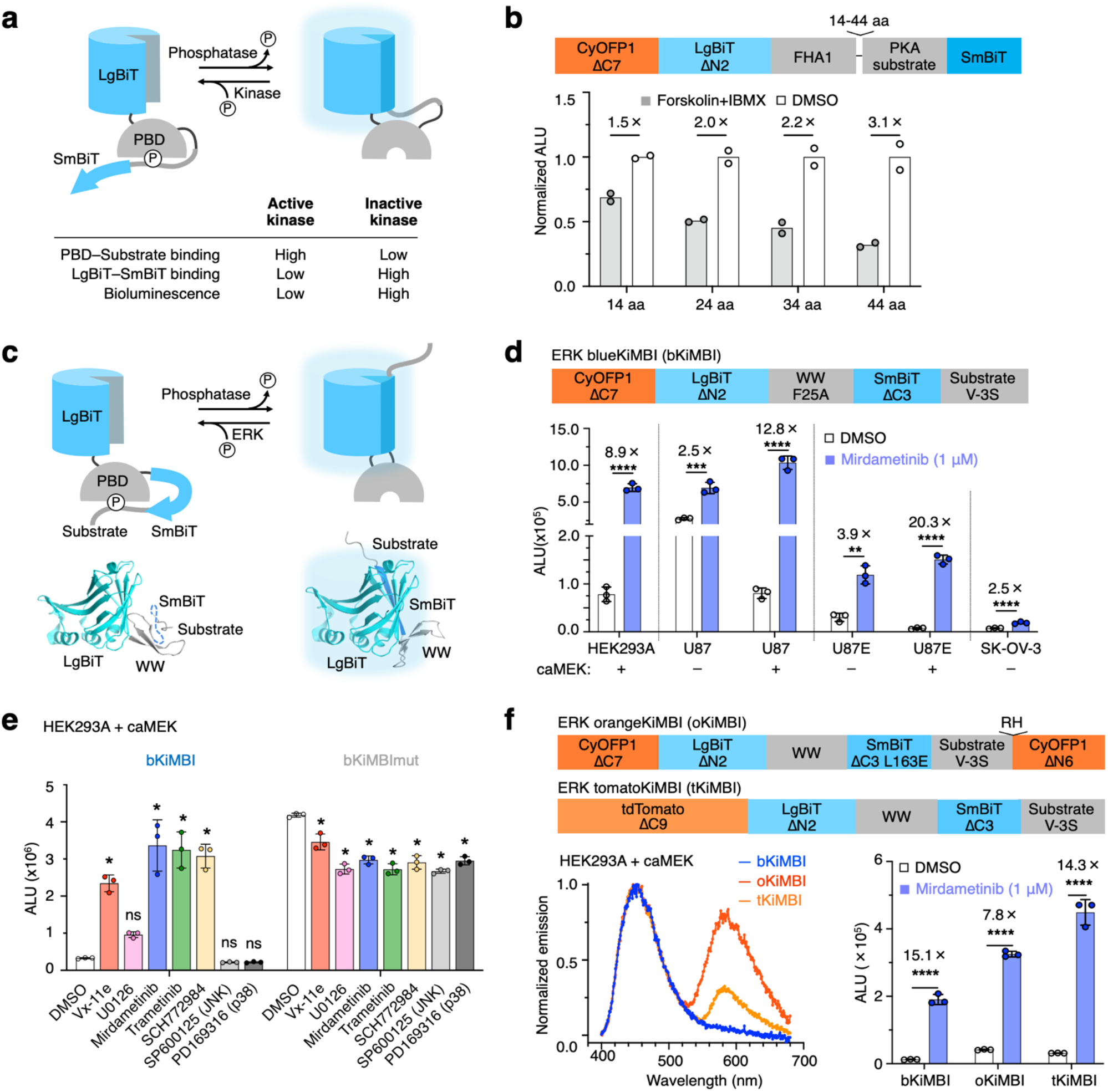
Development and characterization of KiMBIs. **a,**Proposed mechanism of KiMBI. With active kinase, phosphorylated kinase substrate binds to phosphopeptide-binding domain (PBD), outcompeting LgBiT–SmBiT reconstitution (left). Kinase inhibition allows dephosphorylation and LgBiT–SmBiT reconstitution (right). **b**, Bioluminescence of cells expressing putative PKA KiMBIs with different linker lengths between PBD and substrate. All variants are inhibited by PKA activators Fsk and IBMX. **c,**Domain arrangement and model of an ERK KiMBI. **d,**bKiMBI signals with or without caMEK cotransfection or MEK inhibitor mirdametinib. U87E is U87-EGFRvIII. **, p < 0.01; ***, p < 0.001; **** p < 0.0001 by unpaired twotailed Student’s test. Error bars show s.d. **e**, Mean bioluminescence of HEK293A cells co-expressing caMEK and tKiMBI or tKiMBImut negative control in response to inhibitors Vx-11e (ERK, 1 μM), U0126 (MEK, 10 μM), mirdametinib (MEK, 1 μM), trametinib (MEK, 1 μM), SCH772984 (ERK, 1 μM), SP600125 (JNK,10 μM), and PD169316 (p38, 10 μM). One-way ANOVA (p < 0.0001) was followed by Tukey’s posthoc test; ns, p > 0.05; *, p < 0.05. **f,**Top, domain arrangements of orangeKiMBI and tomatoKiMBI. Left, bioluminescence spectra of KiMBI color variants with caMEK cotransfection, normalized to the 450-nm peak. Right, responses of color variants to mirademetinib. ****, p < 0.0001 by unpaired two-tailed Student’s t-test. Error bars show s.d. ALU, arbitrary luminescence units.

We first tested our design using a PKA substrate and a cognate FHA domain^25^. We generated proteins with the topology LgBiT-FHA-linker-substrate-SmBiT, screening linker lengths to allow the phosphorylated substrate to reach its binding site on FHA. Proteins were expressed in mammalian cells, which were then treated with forskolin and IBMX to activate endogenous PKA. All constructs showed lower bioluminescence with PKA activation, with the longest linker (44 amino acids) producing the largest induction (**Fig. 1b**).

Given the importance of the Ras-ERK pathway in human disease, we next aimed to develop a bioluminescent indicator for ERK inhibition. We created and tested fusion proteins of various topologies (**Fig. S1a**) comprising the NanoBiT fragments, an ERK substrate peptide from Cdc25C^26^, and a proline-directed WW phospho-binding domain (WW)^27^ from Pin1. We screened the KiMBI candidates by co-transfection with a constitutively active MEK (caMEK) followed by treatment with the ERK inhibitor Vx-11e. One topology produced an increase in luminescence (Δ*L*/*L*) of ~65% when treated with 1 μM Vx-11e (**Fig. S1a**). In this top responder (LgBiT-WW-SmBiT-substrate), binding of pSub to WW is expected to force the SmBiT fragment to adopt an unfavorable conformation for its complementation with the LgBiT, while substrate dephosphorylation should allow LgBiT-SmBiT complementation and luciferase activity restoration (**Fig. 1c**). These results demonstrate that bioluminescent indicators of kinase inhibition can be engineered on the general concept of PBD-pSub binding competing with LgBiT-SmBiT binding.

### Optimization and characterization of the ERK KiMBI prototype

We next explored optimizing the performance of ERK KiMBI. Based on the proposed mechanism of action, we reasoned that disfavoring LgBiT-SmBiT complementation or enhancing WW-phosphosubstrate binding could lower the background signals when ERK is active, thus improving the signal fold change upon inhibitor treatment. By screening truncated or mutated SmBiT variants, we found truncation of 3 amino acids (aa) from the SmBiT C-terminus (CT) improved the response to ~3-fold (ΔC3, **Fig. S1b**). We then further improved ΔC3 by introducing a second ERK phosphorylation site at position −3 within the Cdc25C substrate sequence to increase WW-pSub avidity. This change improved the induction to 8.8-fold (ΔC3 V–3S, **Fig. S1c**). Substitution of either the additional (serine, position −3) or original (threonine, position 0, **Fig. S1c**) phosphorylation site with alanine notably decreased the drug response, suggesting that both sites are phosphorylated by ERK and recognized by the WW domain (**Fig. S1d**). Finally, we further strengthened WW-phosphosubstrate binding by introducing an F25A mutation in the WW domain^28^ (**Fig. S1e**). The resulting construct showed a >10-fold response by Vx-11e, and was named ERK blue KiMBI (bKiMBI, **Fig. S1e**).

We validated bKiMBI function in cancer cell lines with hyperactive ERK, choosing U87 (human glioma), U87-EGFRvIII (U87 expressing epidermal growth factor receptor variant III, which further activates ERK), and SK-OV-3 (human ovarian cancer). As expected, bKiMBI showed significant signal increases after ERK inhibition in all tested cell lines. U87-EGFRvIII exhibited a lower basal KiMBI signal than U87, consistent with its higher basal ERK activity due to the upstream constitutive activator EGFRvIII (**Fig. 1d**). Endogenous ERK activity in these cell lines appeared to be only partially activated, as co-expressing caMEK further suppressed basal KiMBI signal. Also, we found that ERK bKiMBI was activated by various MEK or ERK inhibitors, but not by any JNK or p38 inhibitors (**Fig. 1e**). This provides additional evidence that KiMBI responds positively to ERK inhibition, and argues against inhibitors directly binding KiMBIs to enhance NanoLuc assembly or catalysis. As a control, we also tested the effects of kinase inhibitors on a phosphosite mutant of ERK bKiMBI (bKiMBImut, A-A in **Fig. S1e**); this revealed no response to kinase inhibition.

Next, to red-shift KiMBI emission for better tissue penetration *in vivo,* we explored fusing fluorescent proteins to bKiMBI to allow resonance energy transfer (RET). Although constructs in the early rounds of engineering included a CyOFP1^29^ at their N-terminus, there was no detectable RET between the reconstituted NanoBiT and CyOFP1 (**Fig. S1f,g**). Following the example of Antares luciferase^29^, we tried fusing a second CyOFP1 domain to the C-terminus of the ΔC3 V–3S KiMBI precursor. Only the additional fusion of a C-terminal CyOFP1 with a 2-aa linker improved RET efficiency, but it also increased background luminescence and diminished the response to ERK inhibition (C6, **Fig. S1f,g**). To rescue the response, we weakened the ability of SmBiT to bind LgBiT by performing alanine scanning followed by further mutagenesis, resulting in an optimized L163E mutation (**Fig. S1h, Table S1** and **Table S2**). The resulting construct, named orange KiMBI (oKiMBI), maintained high RET while restoring most of the inducibility of bKiMBI (**Fig. 1f**).

We also tested other potential RET acceptors fused to the N-terminus of LgBiT in place of CyOFP in bKiMBI (**Table S3**). Fusion with tdTomato allowed a high amount of RET and even better inducibility by the inhibitor (**Fig. 1f** and **Table S3**), and was named tomato KiMBI (tKiMBI). Interestingly, the F25A mutation that had improved bKiMBI responses did not improve oKiMBI or tKiMBI responses, so Phe-25 was retained in these variants (**Fig. S1i**). oKiMBI and tKiMBI were activated by all MEK or ERK inhibitors tested, but not by JNK or p38 inhibitors (**Fig. S1j,k**), reconfirming specificity for ERK inhibitors.

We finally attempted to further improve tKiMBI responsivity by duplicating the WW domain and substrate peptide to add avidity and reduce signal in the kinase-active state (**Fig. S2a,b**). Relative responses did indeed improve (**Fig. S2c**), but maximal brightness and induction kinetics of the sensor were impaired (**Fig. S2c,d**). Thus, we chose the original design of tKiMBI for further application in vivo.

### Molecular imaging of ERK inhibition in a subcutaneous tumor xenograft model

To report ERK kinase inhibition in human tumor xenografts in mice, we first generated U87-EGFRvIII stable cell lines expressing tKiMBI. U87-EGFRvIII cells were transduced with lentiviruses expressing tKiMBI-T2A-AkaLuc, where AkaLuc allows kinase-independent tracking of tumor location and size (**Fig. 2a** and **Fig. S3a**). Transduced cells were sorted by fluorescence-activated cell sorting (FACS) and selected by antibiotics to generate polyclonal stable cell lines. A control reporter cell line expressing the phosphosite mutant of tKiMBI (tKiMBImut) was generated in parallel, and the expression of the indicators was confirmed by fluorescence microscopy (**Fig. S3b**).

**Fig. 2.**
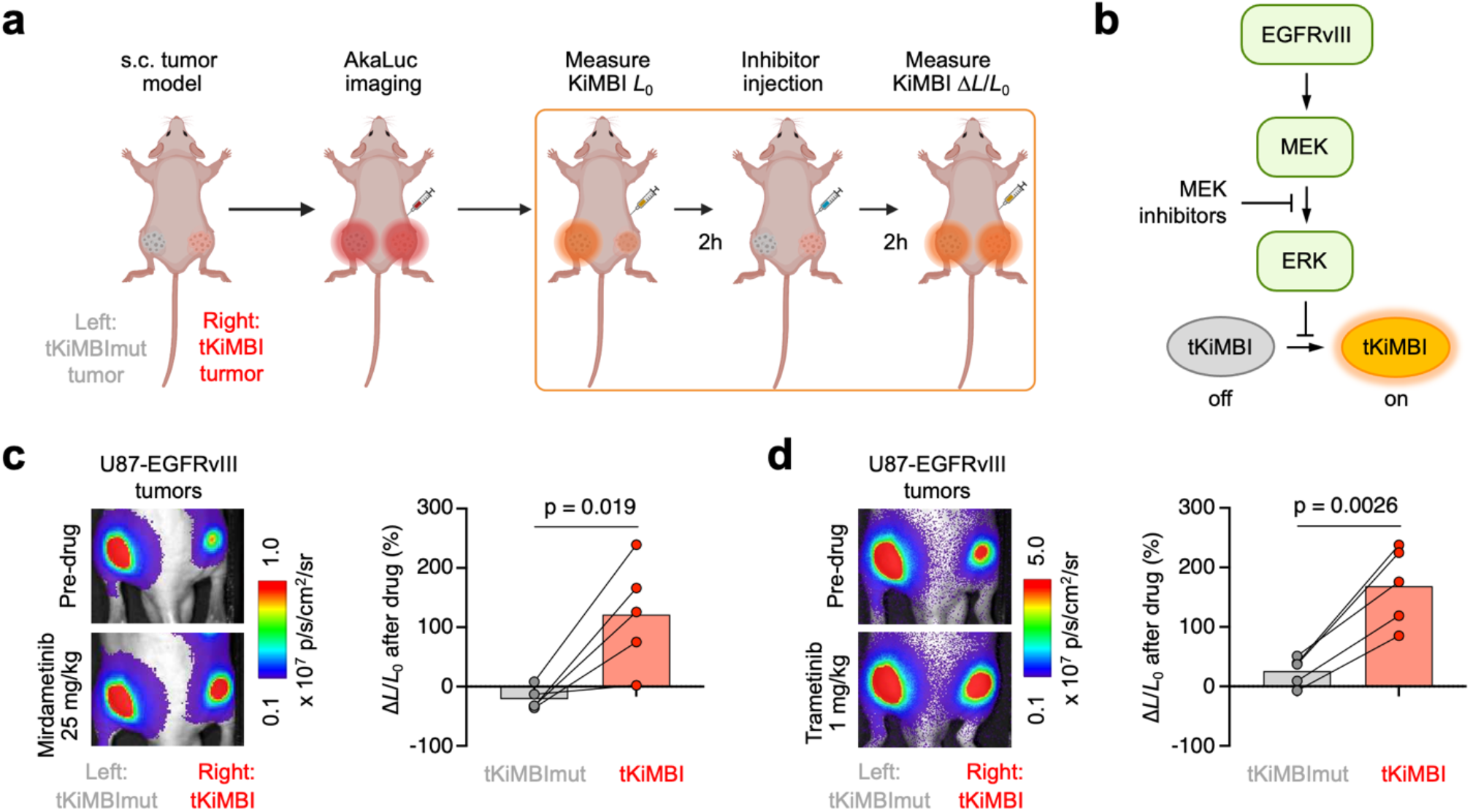
Molecular imaging of Ras-ERK pathway inhibition in a subcutaneous tumor model. **a,**Scheme of the experimental design. U87-EGFRvIII reporter cells expressing tKiMBI (right) and tKiMBImut (left) were implanted subcutaneously to establish tumors in J:NU mice. AkaLuc imaging with AkaLumine injection was used to monitor tumor growth. To visualize ERK inhibition, the indicators were imaged with FFz injection, 2 h before and 2 h after treatment with MEK-ERK inhibitors. All injections were performed intraperitoneally (i.p.). **b**, Scheme of ERK signaling pathway and MEK inhibition. **c-d,**U87-EGFRvIII tumor-bearing mice were sequentially treated (two days apart) with MEK inhibitors mirdametinib (**c**) and trametinib (**d**) and were imaged with FFz injection 2 h after inhibitor injection. Left, representative bioluminescent images collected before and after inhibitor treatment. Right, the percentage change of bioluminescence signals collected before and after inhibitor treatment. Each line represents an individual mouse. P values, paired two-tailed Student’s t-test. L, luminescence.

These U87-EGFRvIII cells stably expressing tKiMBI exhibited ~4-fold bioluminescence increases upon treatment with mirdametinib (MEK inhibitor, previously PD0325901), trametinib (MEK inhibitor, GSK1120212), or SCH772984 (ERK inhibitor) (**Fig. S3c**). In contrast, the tKiMBImut-expressing cells did not show induction of bioluminescence by these inhibitors. Thus, stably expressed tKiMBI responded to MEK or ERK inhibitors similarly to transiently expressed tKiMBI.

We then assessed the ability of tKiMBI to report ERK pathway inhibition in tumors in vivo by the clinically approved MEK inhibitors mirdametinib and trametinib (**Fig. 2b**, **Fig. S4a**). Mice were injected subcutaneously with 10^6^ tKiMBI-expressing cells on one side and, to control for differences in substrate delivery between injections, tKiMBImut-expressing reporter cells on the other side (**Fig. 2a**). One week after tumor cells implantation, sizes of the engrafted tumors were quantified by measuring peak AkaLuc bioluminescence after injecting AkaLumine (**Fig. S4b**). We then imaged tKiMBI bioluminescence before or 2 h after injection of mirdametinib by administering the NanoLuc substrate fluorofurimazine (FFz)^17^. We saw a clear signal increase after mirdametinib treatment in the tKiMBI-expressing tumor, but not in the tKiMBImut-expressing tumor (**Fig. 2c, Fig. S4c,d**). On average, tKiMBI responded to mirdametinib with a 120% increase in brightness, significantly different from the –20% signal change of tKiMBImut (**Fig. 2c**). This result illustrates the ability of tKiMBI to report ERK inhibition within tumor cells in vivo.

An advantage of non-invasive tKiMBI imaging is that multiple kinase inhibitor candidates can be tested in one mouse by repeating the imaging procedure. For example, after waiting 2 days for injected mirdametinib to clear, a second test on the same group of mice with trametinib also revealed successful ERK pathway inhibition (**Fig. 2d, Fig. S4e,f**). Thus, tKiMBI allows multiple kinase inhibitors to be tested and compared in a single mouse.

### Reporting BBB permeability of kinase inhibitors by expressing KiMBI in the brain

For kinase inhibitors to effectively treat brain tumors, BBB permeability is crucial. Gliomas in the central nervous system can be either diffusively infiltrating without BBB breakdown, or can create large masses with central regions of BBB remodeling, forming a blood-tumor-barrier (BTB) that is typically more permeable^30^. However, even in gliomas with permeable BTBs, cancer cells migrate centrifugally into regions of brain parenchyma with intact BBB^31^. Indeed, most gliomas recur after the removal of the visible tumor due to the difficulty of completely removing these infiltrative tumor cells. Kinase inhibitors that can penetrate the BBB thus are more likely to be effective for diffusing tumors and for preventing the recurrence of gliomas after surgery.

Having validated tKiMBI’s ability to respond to MEK inhibitors in vivo, we next asked whether tKiMBI expressed in the brain could provide a rapid method for assessing BBB permeability of kinase inhibitors. To report the BBB permeability of Ras-ERK pathway inhibitors, we used AAV to co-express tKiMBI and caMEK in mouse brains, where caMEK serves to raise the baseline level of ERK activity (**Fig. 3a**). Since the stoichiometric ratio between caMEK and tKiMBI may affect the performance of the sensor, we compared bicistronic reporter genes with opposite arrangements, caMEK-T2A-tKiMBI (A1) and tKiMBI-T2A-caMEK (A2) in cell-based bioluminescence assays (**Fig. S5a**). When transiently transfected in HEK293A cells, both A1 and A2 reporter genes produced an approximately 4-fold increase in tKiMBI signals upon mirdametinib treatment (**Fig. S5b**). A2 was chosen for further characterization using more MEK/ERK inhibitors, and demonstrated satisfactory responsivity (**Fig. S5c**). After being packaged into AAV vectors, the reporter (tKiMBI-T2A-caMEK) or its negative control (tKiMBI_mut_-T2A-caMEK) was expressed in mouse striatum by AAV transduction (**Fig. 3a**).

**Fig. 3.**
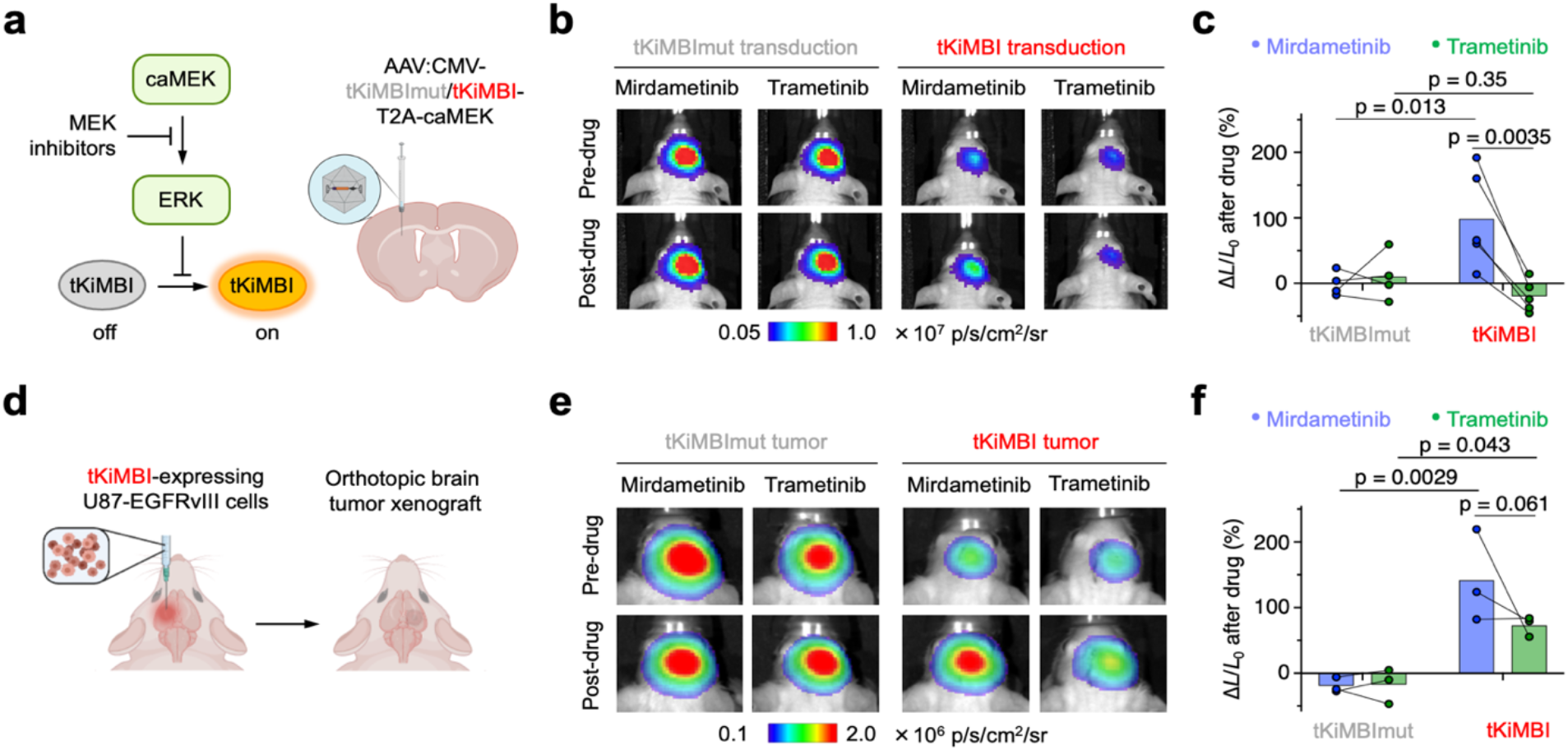
Molecular imaging of ERK inhibition by MEK inhibitors in mouse brain. **a,**Scheme of AAV infection for KiMBI expression in the mouse striatum and pathway schematic with position of MEK inhibitors. **b**, 4 weeks after AAV infection in the striatum, tKiMBI- or tKiMBImut-expressing J:NU mice were sequentially treated (two days apart) with mirdametinib or trametinib and were imaged with CFz injection. **b,**Representative bioluminescence images collected 2 h before and 2 h after inhibitor treatment. **c,**The percentage change of bioluminescence signals collected before and after inhibitor treatment. P values, one-way ANOVA analysis (p = 0.006) was followed by Holm-Sidak’s posthoc test. Each line represents an individual mouse. **d**, Design of the tKiMBI-expressing brain tumor xenograft model. **e**, J:NU Mice with tKiMBI- or tKiMBImut-expressing U87-EGFRvIII tumor engrafted in the striatum were sequentially treated (two days apart) with mirdametinib or trametinib and imaged with CFz injection. Representative bioluminescence images were collected before and after inhibitor treatment. **f,**The percentage change of bioluminescence signals collected before and after inhibitor treatment. P values, one-way ANOVA analysis (p = 0.0025) was followed by Holm-Sidak’s posthoc test. Each line represents an individual mouse.

We then tested whether tKiMBI could report the efficacy of MEK inhibitors in the brain, using mirdametinib and trametinib as reference compounds. Mirdametinib accumulates in brain tissue and blocks ERK phosphorylation in the brain after peripheral administration^32, 33^. In contrast, pharmacokinetic and pharmacodynamic analyses did not detect trametinib presence or ERK inhibition in the brain after peripheral injection^32–35^. We then measured tKiMBI responses upon treatment with mirdametinib or trametinib at doses effective against extracranial tumors^34, 36^, using recently reported brain-optimized NanoLuc substrates (Su et al., in press). The negative control tKiMBI_mut_ did not respond to either drug (**Fig. 3b, Fig. S5d-g**), confirming that the drugs do not directly inhibit NanoLuc or alter substrate levels in the brain. As expected, mirdametinib robustly induced a significant increase in tKiMBI signal, confirming its ability to inhibit the Ras-ERK pathway in mouse brain (**Fig. 3b, Fig. S5d,e**). In contrast, trametinib produced no response from tKiMBI (**Fig. 3b, Fig. S5f,g**), consistent with its previously characterized BBB impermeability. By using the same mice to assess mirdametinib and trametinib, we could conclude trametinib was less BBB-permeable with high confidence (**Fig. 3c**). These results demonstrate that tKiMBI expressed by AAV in mouse brains reliably reports BBB permeability of inhibitors targeting the Ras-ERK pathway.

We next tested whether KiMBI can report ERK inhibition in an orthopic xenograft model of brain cancer (**Fig. 3d**). Specifically, we implanted 3 × 10^4^ tKiMBI- or tKiMBI_mut_-expressing U87-EGFRvIII cells into the striatum and assessed the response of tKiMBI to peripherally injected mirdametinib or tramertinib. As U87-EGFRvIII brain xenografts are known to produce a permeable BTB in mice^30^, we hypothesized that both mirdametinib and tramertinib would be able to enter the KiMBI-expressing tumor cells in the brain. Indeed, we observed ~180% and ~130% increases in tKiMBI signal after mirdametinib and trametinib administration, respectively (**Fig. 3e,f, Fig. S6**).

As the provenance of U87 cells distributed by North American repositories is unknown^37^, we also assessed KiMBI responses in a second widely used human glioblastoma line, LN-229. Cultured LN-229 cells stably expressing tKiMBI exhibited a roughly 1.6-fold increase in bioluminescence after treatment of Ras-ERK pathway inhibitors (**Fig. S7a–c**), demonstrating some baseline ERK activity, albeit at lower levels than in U87-EGFRvIII cells. We implanted 3 × 10^4^ LN-229 cells expressing tKiMBI or tKiMBI_mut_ into the mouse striatum, and then performed bioluminescent imaging one week later. Treatment with either mirdametinib or trametinib resulted in >3-fold signal increases from tKiMBI, suggesting that LN-229 xenografts also produce a leaky BTB (**Fig. S7d-g**). Taken together, these results demonstrate that the BTB is leaky in commonly used orthotopic xenograft mouse models of glioma.

### Non-invasive characterization of drug pharmacodynamics in the body and brain

Noninvasive evaluation of kinase inhibition over time in target tissues, i.e. pharmacodynamics, would be extremely desirable for assessing the optimal concentration and dosing intervals of drug candidates. For cancer indications in the brain, comparing pharmacodynamics in the presence of an intact BBB versus a leaky BTB could help rule out any efficacy differences between bulk tumor and tumor margin^31^. The brain-optimized NanoLuc substrate demonstrates rapid signal decay with a half-life of ~8 min, allowing repeated imaging with ≥1-h intervals (Su et al., in press). We thus performed repeated bioluminescence imaging of ERK KiMBI to track pharmacodynamics of the BBB-permeable MEK inhibitor mirdametinib. Specifically, we tested pharmacodynamics in tumor xenograft models outside and inside the brain, as well as intact brain parenchyma (**Fig. 4a**).

**Fig. 4.**
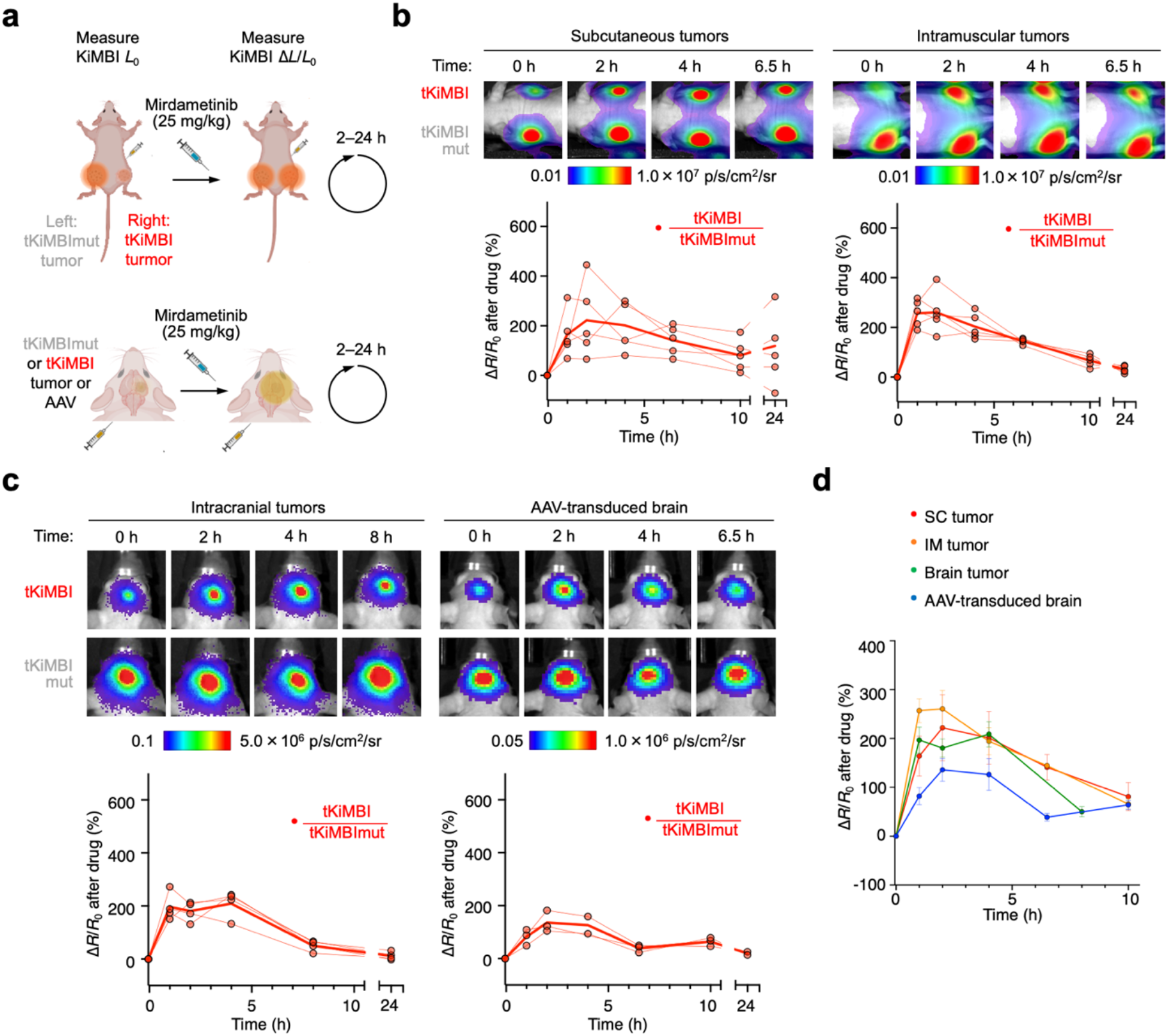
Molecular imaging of pharmacodynamics of ERK inhibition by mirdametinib in multiple disease models. **a,**Design of experiments to measure ERK inhibitor pharmacodynamics. Initial bioluminescence L_0_ from reporter-expressing cells was measured with CFz injection 2 h before inhibitor treatment. After inhibitor administration, CFz was reinjected and bioluminescence L measured at various time points. **b,**Mirdametinib pharmacodynamics in extracranial tumor models. Top, representative bioluminescence images collected at indicated time points after inhibitor. Bottom, pharmacodynamics timecourse of tKiMBI/tKiMBImut ratio changes (ΔR/R_0_). **c,**Mirdametinib pharmacodynamics in tKiMBi-expressing intracranial xenografts or brains transduced with reporter and caMEK. **d,**Summary of pharmacodynamic time-courses of ERK inhbition by mirdametinib in different tissues.

Treatment with mirdametinib could conceivably alter tumor bloodflow or reporter expression at extended timepoints which could affect tKiMBI brightness, but these indirect mechanisms should affect tKiMBImut as well. Thus for intramuscular or subcutaneous tumors, we implanted tKiMBI-expressing and tKiMBImut-expressing cells on opposite sides in the same mice (**Fig. S8a**) and normalized the fold change in tKiMBI intensity at each time with the tKiMBImut measurement from the contralateral tumor. For cranial tumors and AAV-transduced brain, we implanted tKiMBImut-expressing cells or injected tKiMBImut-expressing AAV in a parallel control cohort (**Fig. S8b**) and normalized the fold change in tKiMBI intensity at each time with the mean tKiMBImut measurement from the simultaneously imaged control cohort (**Fig. S8c**).

In intramuscular or subcutaneous tumors, tKiMBI signals reached a peak response 2 h after peripheral administration of 25 mg/kg mirdametinib, then decayed with a half-life of approximately 6 h (**Fig. 4b**). Peak response amplitudes were ~200% after correction by the tKiMBImut signal. Similar results were observed for tKiMBI in intracranial tumors (**Fig. 4c,d**). In tKiMBI-transduced brain tissue, kinetics were not discernably different in either time to peak or rate of decay (**Fig. 4c,d**). These results suggest that clearance rates were not slower in the brain than in plasma for mirdametinib; i.e. there was no discernable depot effect in the brain. Taken together, these results demonstrate that longitudinal tKiMBI imaging can reveal kinase inhibitor pharmacodynamics in bulk tumors or in normal tissue noninvasively.

### Identification of an ERK inhibitor with activity in the brain

Given the success of AAV-transduced tKiMBI in reporting the BBB permeability of well-characterized MEK inhibitors, we finally asked if tKiMBI can identify ERK inhibitors with high efficacy in the brain. We found that no ERK inhibitors have been unambiguously identified as BBB-penetrant, and reports on brain concentrations have not been published for most of them. We selected KO-947, ulixertinib, and temuterkib (LY3214996) for testing with ERK tKIMBI due to their promising performance in pre-clinical trials for treating extracranial tumors (**Fig. S9a**)^38^.

We first confirmed the in vivo efficacy of KO-947, ulixertinib, and temuterkib using a tKiMBI-expressing intramuscular xenograft mouse model (**Fig. S9a**). As expected, all three ERK inhibitors induced a signal increase in tKiMBI-expressing tumors (**Fig. S9b-d**). Next, using the tKiMBI brain AAV model where caMEK and tKiMBI are coexpressed (**Fig. 5a**), we examined the efficacy of the ERK inhibitors in mouse brains. Despite promising efficacy in intramuscular models, treatment with KO-947 and ulixertinib did not induce any significant response of tKiMBI (**Fig. 5b-c, Fig. S10a-b**). These results demonstrated that KO-947 and ulixertinib cannot efficiently penetrate the BBB and inhibit the ERK activity in brain tissues. However, we found temuterkib to be efficacious in the AAV-transduced brain model (**Fig. 5d, Fig. S10c**), demonstrating the BBB permeability of temuterkib. Our findings suggest that temuterkib is a more promising candidate ERK inhibitor than KO-947 and ulixertinib for brain indications.

**Fig. 5.**
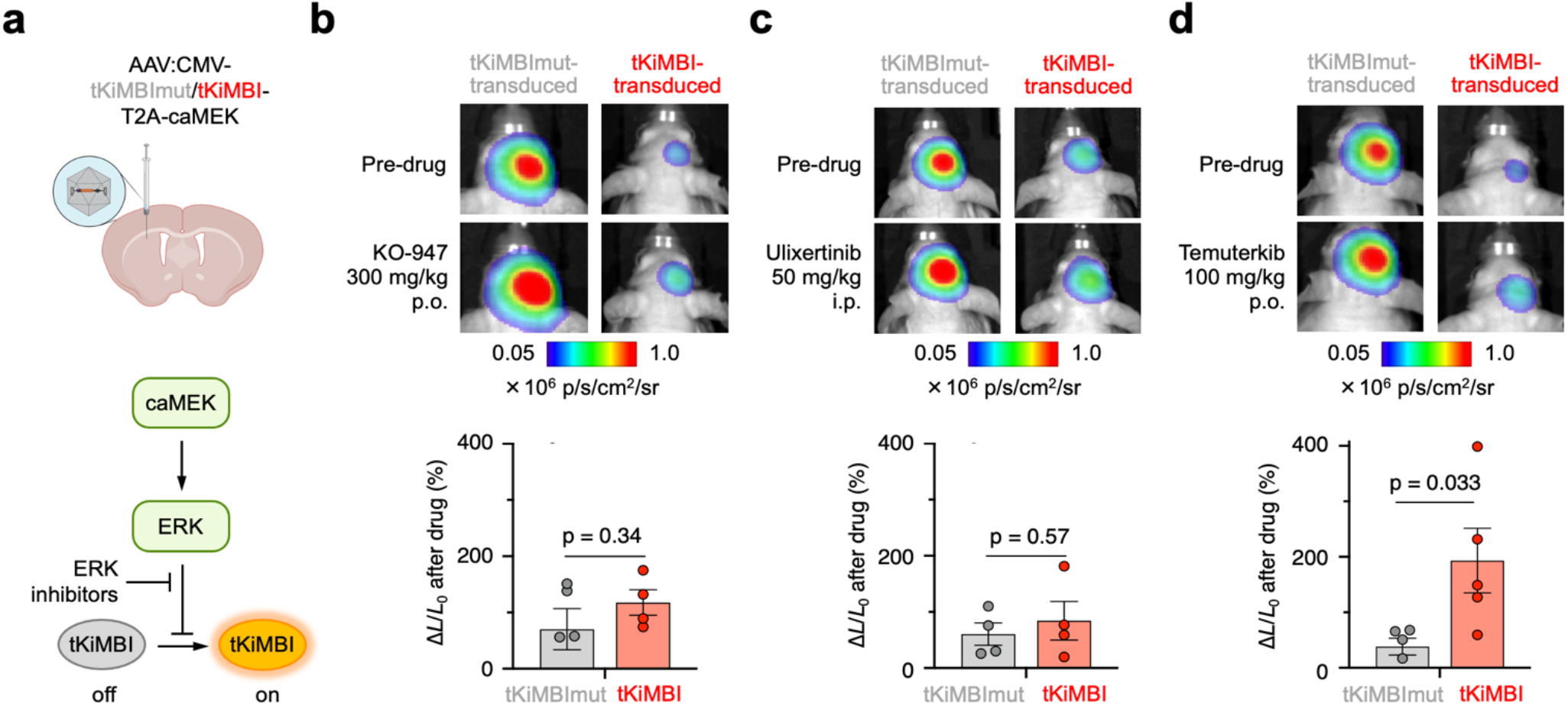
Molecular imaging of ERK inhibitors in AAV-infected mouse brain. **a**, Pathway schematic with position of ERK inhibitors. **b-d**, After AAV infection in the striatum, tKiMBI- or tKiMBImut-expressing J:NU mice were imaged 2 h after ERK inhibitor treatment. Above, representative bioluminescence images collected before and after each ERK inhibitor treatment. Below, percentage change of bioluminescence signals collected before and after inhibitor treatment. P values, unpaired two-tailed Student’s t-test.

Finally, we asked whether our findings could be reliably predicted in silico. Algorithms have been developed to predict BBB permeability of drugs based on their physicochemical properties, among which the “BBB score” is the most recent, outperforming earlier approaches^39^. Using the “BBB score calculator” on KO-947, ulixertinib, and temuterkib, we found KO-947 to have the highest BBB score and temuterkib the lowest (**Table S4**). This is contrary to our empirical findings that temuterkib but not KO-947 is effective in the brain. Thus, while the physiochemical properties of candidate drugs certainly affect BBB penetration, the ability of compounds to accumulate in the brain is apparently still difficult to predict, underscoring the utility of empirical approaches such as KiMBI imaging.

## Discussion

In this study, we addressed the need for non-invasive imaging of kinase inhibition in the brain by creating kinase-modulated bioluminescent indicators (KiMBIs) that use optimized luciferase substrates. We find that an ERK KiMBI reports Ras-ERK pathway inhibition in tumors in mice, and can differentiate between BBB-permeable and BBB-impermeable inhibitors. We used ERK KiMBI to demonstrate BTB disruption in xenografts and track MEK inhibitor pharmacodynamics in an orthotopic glioma model and in normal brain tissue. Finally, we used ERK KiMBI to evaluate the intracranial activity of ERK inhibitors, identifying temuterkib to be suitable for use in the brain, despite algorithmic predictions of poor BBB permeability. The KiMBI technology described here provides a rapid and inexpensive method to empirically assess kinase inhibition in target tissues in living mice.

Our study demonstrates the advantages of non-invasive imaging over earlier methods for empirical evaluation of kinase inhibitors in the brain. Traditional methods for characterizing drugs in animal models require tissue collection and biochemical analysis, which are time-, labor-, and resource-consuming. In particular, such experiments require large numbers of animals, due to experimental variation between animals coupled with the inability of each animal to contribute more than one time point. In contrast, bioluminescence imaging is rapid and inexpensive, involving only injection of the luciferase substrate, anesthesia, and imaging. More importantly, by being noninvasive, KiMBI imaging allows a single experimental subject to be reused multiple times.

For example, a small set of KiMBI-expressing mice can be used to report on multiple drugs (**Fig. 2**), or to test different doses for a drug. Likewise, a small set of animals can report drug efficacy over time, as exemplified by our pharmacodynamic study on mirdametinib (**Fig. 4**). There, genetic encoding and the timelapse nature of observation minimizes experimental noise from different tumor sizes. In traditional pharmacodynamics experiments, different tumor sizes would be expected to introduce variability in kinase activity measurements, as tumor size could influence pathway activity, protein concentration, or the degree of contamination of tumor sample by non-cancerous stroma. However, in xenograft models, KiMBI would only be expressed in the tumor cells, and its response to drug can be normalized to its pretreatment intensity. Thus, our mirdametinib experiment required only 4 mice to obtain 7 time points in the extracranial xenograft models, whereas traditional pharmacodynamic measurements would have required at least 28 mice, and likely more to compensate for variability from using different mice to construct a time course. Indeed, we were unable to find examples in the literature of time-lapse phamacodynamic profiling of mirdametinib; the sole example of pharmacodynamics actually only investigated the effects of two doses at a single timepoint^40^. KiMBI pharmacodynamic reporting can be useful for avoiding underdosing in mouse efficacy testing or for screening candidate inhibitors for prolonged pharmacodynamics.

Finally, while we used caMEK and ERK KiMBI to assess the efficacy of MEK and ERK inhibitors in the brain, it should also be possible to use Ras-ERK pathway activators that are further upstream. This would allow comparison in the same mice of the pharmacodynamic effects on pathway output (ERK activity) by inhibitors targeting the various levels of the pathway, which could have different effects due to feedback regulation. By increasing the pharmacodynamic information obtainable from each animal subject by an order of magnitude, KiMBI reporters should facilitate new kinase inhibitor discovery and dose optimization.

Interestingly, to our knowledge, no bioluminescent reporter for the Ras-Raf-MEK-ERK pathway of any design has been reported. This is remarkable considering that proteins in this pathway are major drug targets in cancer, and that FLuc-based reporters have been reported for Akt, c-Met, EGFR, and ATM^15, 16, 41^. The development of ERK KiMBI thus also fills a previously unmet need for non-invasive reporting of Ras-ERK pathway activity in mammals.

Another conclusion of this study is that xenografts in the mouse brain may falsely report kinase inhibitor efficacy due to BTB leakiness. Specifically, we observed the BBB-impermeable trametinib still activating ERK KiMBI in U87-EGFRvIII and LN-229 xenografts in the brain. However, drugs with good BBB permeability are still required for maximally effective treatment of glioblastoma patients, as low-grade gliomas and a portion of progressive glioblastomas exhibit an intact BBB^42^. In addition, human xenografts in the mouse brain may exhibit more BBB disruption than the human tumors they are intended to model^43^. Evaluating candidate kinase inhibitors by only brain tumor shrinkage in mice may thus produce false-positive results, in addition to requiring large sample sizes for each compound and dosing regimen due to high variability in growth rates between individual xenografts. In contrast, comparing drug responses of KiMBI transduced into cells inside and outside the brain should provide a reliable non-invasive measure of BBB permeability.

A common question regarding bioluminescent indicators is how their use case differs from fluorescent indicators. Kinase indicators based on fluorescent proteins are widely used for live-cell experiments, where their high photon flux enables visualization of kinase activity with spatial and temporal resolutions on the micron and millisecond scales respectively^1^. However, the excitation light required for fluorescence imaging is difficult to deliver uniformly through tissue. Thus, fluorescent indicators can only be used in living mammals by inserting lenses of fibers to access specific sites of interest^44^. This is invasive, severely restricts the field of view, and adds to experimental variability between animals. In contrast to fluorescence, bioluminescence from luciferase proteins is suitable for non-invasive imaging at millimeter and second scales. The absence of excitation light and autofluorescence essentially eliminate background, allowing detection of the relatively low photonic output of luciferase reporters from deep locations without implanted optical elements, with sensitivity limited only by camera dark current and read noise^14^.

NanoLuc-based KiMBIs differ from earlier kinase reporters based on FLuc in two important ways. First, Nanoluc-RFP fusions with the substrate cephalofurimazine produce an order of magnitude more light in the brain than FLuc with either D-luciferin or CycLuc1 (Su et al., in press). The higher achievable brightness of NanoLuc-based reporters should allow drug effects to be detected at lower levels of reporter expression or with fewer cells than FLuc-based reporters, which is less likely to interfere with normal biological processes. Second, because FLuc is ATP-dependent, it has the potential to generate spurious signals if kinase inhibition, or off-target drug activity, alters cellular energy states. Indeed, FLuc alone serves as a reporter of ATP levels in cells^45^. In contrast, as NanoLuc is ATP-independent, KiMBIs can report kinase inhibition independently of ATP.

The modular design of KiMBIs should facilitate generalization to other kinases of therapeutic interest. The engineering of PKA and ERK KiMBIs demonstrated here shows that constructing a reporter to respond positively to kinase inhibition may require different ordering of elements (LgBiT, SmBiT, substrate, and PBD) for different kinases. Presumably, this is due to the distinct orientations that phosphosubstrates can interact with different PBDs relative to PBD termini (FHA for PKA, WW for ERK). We expect other PBDs such as 14-3-3 or SH2 domains could replace the WW domain in ERK KiMBI to develop indicators for a broader range of kinases. As a simple screen of possible topologies with PBD and/or substrate sequences linking LgBiT and SmBiT fragments was sufficient to obtain KiMBIs for PKA and ERK, we expect KiMBIs incorporating other PBDs can also be obtained in this manner.

In summary, we have developed a kinase-modulated bioluminescent indicator (KiMBI) to visualize ERK inhibition by small-molecule drugs in tissues of living mice. Our results demonstrate that non-invasive KiMBI imaging can provide valuable information on target engagement by kinase inhibitors in living subjects, including BBB penetration and pharmacodynamics of drug activity. ERK KiMBI thus provides a rapid method to assess inhibition of the Ras-ERK pathway in the body or the brain, and suggests a general strategy to greatly accelerate the discovery of effective small-molecule inhibitors for other kinases as well.

## Methods

### Chemicals

Drugs used in the paper include ERK inhibitor Vx-11e (Selleck Chemicals), SCH772984 (Selleck Chemicals), KO-947 (MedChem Express), Ulixertinib (Selleck Chemicals) and LY3214996 (Selleck Chemicals); MEK inhibitors U0126 (Tocris), PD0325901 (Selleck Chemicals, ApexBio) and GSK1120212 (Selleck Chemicals; ApexBio), JNK inhibitor SP600125 (Selleck Chemicals), and p38 inhibitor PD169316 (ApexBio). Antibiotics used in cell culture include Blasticidin (Invivogen), Geneticin (Invitrogen), penicillin/streptomycin (Gemini Bio), ciprofloxacin (Sigma-Aldrich), piperacillin (Sigma-Aldrich).

### Molecular Cloning

DNA primers for molecular cloning were synthesized by Intergrated DNA Technologies. WW domain was PCR amplified from pcDNA3-ERK-SPARK, a gift from Xiaokun Shu (Addgene plasmid # 106921). PCR amplification was typically conducted with ~20 overlapping DNA base primers and the PrimeSTAR HS DNA Polymerase (Clontech) or the Phusion Flash High-Fidelity PCR Master Mix (Thermo Scientific). DNA fragments were spliced by overlap-extension PCR with ~20 overlapping DNA sequences by Phusion Flash High-Fidelity PCR Master Mix. Molecular cloning was typically carried out using infusion HD cloning kit (Takara Bio) with a plasmid vector linearized by restriction enzymes and an assembled DNA insert with ~20 overlapping DNA sequences. Constructed plasmids were verified using Sanger sequencing by Elim Biopharm. The KiMBI constructs were assembled in pcDNA3.1 vector (Addgene) behind the CAG promoter. The best KiMBI sensors (bKiMBI, oKiMBI, and tKiMBI) were then cloned into the lentiviral vector pLL3.7m^46^ or the AAV packaging vector pAAV under the CMV promoter.

### Cell lines

HEK293A cells (Invitrogen) were cultured at 37 °C with 5% CO_2_ in Dulbecco’s Modified Eagle’s Medium (DMEM) supplemented with 10% fetal bovine serum (FBS), 2 mM L-glutamate, 100 U/mL penicillin and 100 μg/mL streptomycin. U87 (ATCC, No. HTB-14) and U87-EGFRvIII (previously developed in Lin lab by pcDNA3-EGFRvIII transfection and Geneticin selection) cells were cultured at 37 °C with 5% CO_2_ in DMEM supplemented with 10% bovine calf serum (BCS), 2 mM L-glutamate, 100 U/mL penicillin, 100 μg/mL streptomycin, 0.8 mg/mL Geneticin, 10 μg/mL Ciprofloxacin, 10 μg/mL Piperacillin. SK-OV-3 cells (ATCC, No. HTB-77) were cultured at 37 °C with 5% CO_2_ in Roswell Park Memorial Institute (RPMI) 1640 medium supplemented with 10% FBS, 2 mM L-glutamate, 100 U/mL penicillin and 100 μg/mL streptomycin. LN-229 cells (ATCC, No. CRL-2611) were cultured at 37 °C with 5% CO_2_ in DMEM supplemented with 5% FBS, 2 mM L-glutamate, 100 U/mL penicillin and 100 μg/mL streptomycin.

### Virus packaging

For lentivirus packaging, pLL3.7m-based plasmids were purified by PureLink Expi Endotoxin-free Maxi plasmid Purification kit (Invitrogen), and then HEK293T (ATCC, No. CRL-3216) cells at ~70% confluency were transfected with psPAX2, pMD2.G, and pLL3.7m plasmids using CalPhos mammalian transfection kit (Takara Bio). Two days following transfection, viral supernatant was filtered with a 0.45 μm PES filter before using to infect target cells. For AAV packaging, pAAV-CMV-tKiMBI-T2A-caMEK-bGH_pA plasmids were purified by PureLink Expi Endotoxin-free Maxi plasmid Purification kit (Invitrogen), and then were sent to Stanford Gene Vector and Virus Core to produce and titer AAV.DJ vectors.

### Generation of KiMBI-expressing stable tumor cell lines

U87-EGFRvIII and LN229 cells were lentiviral transduced with pLL3.7m-CMV-KiMBI-T2A-AkaLuc-P2A-Blasticidin. Then, 72 h after transfection, cells were dissociated with trypsin and resuspended in DMEM and then changed to PBS. Suspended cells were sorted for the population with medium fluorescence level (20-40%) on a fluorescence-activated cell sorter (FACS), the FACSJazz (BD Biosciences) at Stanford Shared FACS Facility. After cell sorting, the KiMBI-expressing cells were maintained and passaged in corresponding culture medium supplemented with blasticidin (6 μg/mL for both cell lines) to select for stable polyclonal cell lines. Flow cytometry data were analyzed on FlowJo software. The stable KiMBI expression was validated by examination of tdTomato fluorescence in tKiMBI (and its variants) using an EVOS FL Autoimaging system (Thermo Scientific).

### In vitro cell-based bioluminescent assay

Cells were seeded into 96-well Lumitrac plate (Greiner Bio-One) at a density of 1.5 × 10^4^ to 2 × 10^4^ cells per well. After 24h, cells were transiently transfected with KiMBI-expressing plasmids (5 - 10 ng per well) with or without caMEK-expressing plasmids (50 ng per well, empty pcDNA3.1 as negative control to keep total transfected DNA amount consistent) using lipofectamine 3000 (Invitrogen) following manufacturer’s instructions. For the stable polyclonal KiMBI-expressing cell liens, this lipofection step was skipped. At 24h after transfection, the cells were treated with drugs at desired concentration in opti-MEM reduced-serum medium (Thermo Fisher) for 1–1.5 h. Then, for assaying luminescence, Nano-Glo live cell assay system (Promega) was used following manufacturer’s instructions. Time-lapse live luminescence was recorded on Safire-2 microplate reader (Tecan) with 100 ms integration and one read per min for 20 min. Luminescence spectrum was measured on a Varioskan LUX multimode microplate reader (Thermo Scientific) using the spectral scanning protocol with 1 s of integration for each wavelength at 400–680 nm (1 nm per data point).

### General procedures of kinase inhibitor injection and in vivo bioluminescence imaging

All animal procedures complied with USDA and NIH ethical regulations and approved by the Stanford Institutional Animal Care and Use Committee and according to protocols approved by the Administrative Panel on Laboratory Animal Care (APLAC) of Stanford University. For i.p. drug administration in mice, kinase inhibitors (PD0325901, GSK1120212, KO-947, and Ulixertinib) were dissolved in an injectable formulation containing 5% DMSO, 40% PEG-300, 5% Tween-80 (v/v) in water, with each dose consisting of 25 mg/kg PD0325901, 1 or 3 mg/kg GSK11230212, 50 mg/kg KO-947, or 50 or 100 mg/kg Ulixertinib, in a volume of 150–250 μL. For oral (p.o.) administration, KO-947 and LY3214996 were suspended in a vehicle of 0.5% Methylcellulose and 2% Tween-80 in water, with each dose consisting of 300 mg/kg KO-947 or 100 mg/kg LY3214996. To measure KiMBI intensity, mice were injected i.p. with a solution of 1.3 μmol (0.56 mg) of NanoLuc substrate and 6 mg poloxamer-407 (P-407) in 150 μL of Dulbecco Phosphate Buffered Saline (DPBS, without Ca^2+^ or Mg^2+^, no. 21-031-CV, Corning). Substrate was either fluorofurimazine (FFz) or cephalofurimazine (CFz) or compound 6, a fluorinated analogue of CFz (Su et al., in press). Immediately after substrate administration, mice were anesthetized using isoflurane, and images were acquired in an Ami HT optical imaging system (Spectral Instruments) every 1 min for 10 min, with 1–2% isoflurane in air for anesthesia. Unless otherwise stated in the specific procedures, imaging settings were open emission filter, 25-cm field of view, f/1.2 aperture, 2×2 binning, and 30-s exposure time. Images were analyzed in Aura 4.0 software (Spectral Instruments). The above conditions and settings were for most in vivo studies.

### Molecular imaging of ERK inhibition in subcutaneously or intramuscularly implanted cells

For imaging ERK inhibition in subcutaneously (s.c.) or intramuscularly (i.m.) implanted cells, U87-EGFRvIII cells stably expressing KiMBIs were dissociated with trypsin and resuspended in DMEM and then changed to PBS with a density of 1 × 10^7^ to 2 × 10^7^ cells per mL. For s.c. implantation, 4 × 10^5^ or 1 × 10^6^ cells were resuspended in 100 μL FBS-free Opti-MEM containing 50% Matrigel matrix (Corning). For i.m. implantation, 8 ×10^5^ cells were directly injected in DMEM suspension. First, under sterile conditions, 8- to 10-week-old male nude mice (strain J:NU No.007850, Jackson Laboratories) were anesthetized using isoflurane. Cells were subcutaneously injected into the flanks of nude mice. Mice were recovered on heat pads for 30 min while cells were allowed to settle. One week after cell implantation, for visualizing tumor growth, mice were i.p. injected with 1.5 μmol (0.5 mg) of AkaLumine-HCl (Sigma-Aldrich) in 100 μL 0.9% NaCl for imaging (1 × 1 binning, 10-s exposure time). To visualize ERK inhibition, tumor-bearing mice were i.p. injected with FFz (0.9 μmol, with 6 mg P-407) for imaging. Two hours after the first imaging, kinase inhibitors in the injectable formulation were injected i.p., and a second imaging was carried out 2 h post drug administration following the same procedures as the first imaging.

### Molecular imaging of ERK inhibition in KiMBI-encoded AAV infected mouse brain

Under sterile conditions, the 9-week-old male J:NU mice were anesthetized with isofluorane and secured in a stereotaxic frame (RWD Life Science, Shenzhen, China), and a hole of the size of the needle was drilled through the skull. A Hamilton syringe with a 33-gauge needle was inserted at 0.5 mm dorsal and 2.0 mm lateral to the bregma to a depth of 3.1 mm, and after a 2-min wait the needle was pulled back 0.3 mm to allow space for the virus solution. 1.5 μL of AAV vector solution (2 × 10^12^ viral genomes per mL) was injected at a speed of 0.15 μL/min using a syringe pump (KD Scientific, Holliston, MA). The needle was left in place for 3 min after each injection to minimize the upward flow of viral solution after raising the needle. To visualize ERK inhibition, four weeks after AAV infection, mice were i.p. injected with CFz (1.2 μmol, with 6 mg P-407) for imaging (4×4 binning, 60-s exposure time). Two hours after the first imaging, kinase inhibitors in the injectable formulation were injected i.p., and a second imaging was carried out 2 hours post-drug administration following the same procedures as the first imaging.

### Molecular imaging of ERK inhibition in intracranially implanted cells

The stereotaxic injection on the 7- to 9-week-old male J:NU mice were similar to the afore-mentioned AAV infection surgery. A Hamilton syringe with a 26-gauge needle was inserted 0.62 mm dorsal and 1.75 mm lateral to the bregma to a depth of 3.5 mm, and after a 2-min wait the needle was pulled back 0.5 mm to allow space for the cell suspension. Following this, 3 × 10^4^ KiMBI-expressing U87-EGFRvIII or LN-229 stable cells in 1.5 μL of PBS were injected at an injection speed of 0.3 μL/min using the syringe pump. To visualize ERK inhibition in U87-EGFRvIII cell implants, 5 days after cell implantation, tumor-bearing mice were injected i.p. with CFz (0.4 μmol, with 1.8 mg P-407) for imaging. Two hours after the first imaging, kinase inhibitors in the injectable formulation were i.p. injected, and a second imaging was carried out 2 hours post drug administration following the same procedures as the first imaging. To visualize ERK inhibition in LN-229 cell implants, imaging was done 7 days after cell implantation, and 1.05 or 0.7 μmol CFz with 5 or 3.3 mg P-407 was i.p. injected to each mouse for imaging (4×4 binning, 30-s exposure time).

### Time-lapse imaging of ERK inhibition in intracranially implanted cells

One week after tumor cell implantation or two weeks after AAV injection, the tumor-bearing or AAV-infected mice were i.p. injected with FFz or CFz (0.2 μmol, with 0.8 mg P-407) for the initial imaging to establish the signal baseline before drug administration. Two hours after the first imaging, PD0325901 (25 mg/kg) in the injectable formulation was i.p. injected, and the subsequent imaging was carried out at 1, 2, 4, 8, 24 hours or 1, 2, 4, 6.5, 10, 24 hours post drug administration following the same procedures as the initial imaging.

### Statistics

Student’s t-test, one-way analysis of variance (ANOVA) with Tukey’s or Holm-Sidak’s posthoc test were performed in GraphPad Prism version 9.0.0 (Dotmatics).

## Data availability

The main data supporting the findings of this study are available within the article and supplementary information. Additional raw data are available from the corresponding author upon request.

## Supporting Information

Tables S1 (Alanine scanning on SmBiT), S2 (Mutations on SmBiT sites 161 and 163), S3 (Screening of N-terminal fusions of fluorescent proteins) and S4 (Predicted BBB scores of select ERK inhibitors) (PDF).

## Acknowledgements

We thank Lan Liu (Lin laboratory) for expert assistance with animal breeding and Pamelyn Woo (Monje Lab) for training on orthotopic glioma xenografts. We thank Hokyung Kay Chung for generating the U87-EGFRvIII stable line while in the Lin laboratory. We thank Xiang Wu (laboratory of Guosong Hong, Stanford) for providing training and instrument on stereotaxic injection. We thank Stanford Gene Vector and Virus Core for packaging and tittering AAV vectors, and Stanford Shared FACS facility for providing training and instruments for cell sorting. Illustrations were created with Biorender.com. Funding was provided by NIH grants R21NS122055 (M.Z.L.) and R21DA048252 (M.Z.L.), a Stanford Discovery Innovation Award (M.Z.L.), a Stanford Bio-X Seed Grant (M.Z.L. and M.M.), an Agilent Fellowship (Y.W.), a Stanford Bio-X Bowes Graduate Student Fellowship (Y.W.), NSF I-Corps Grant 2106025 (L.N., M.Z.L.), and a Stanford Bio-X Visiting Scholar Fellowship associated with the Novo Nordisk Foundation grant NNF18OC0031816 (M.W.).

## Author contributions

Y.S., Y.W., L.N., M.W., and M.Z.L. designed experiments. Y.S. and Y.W. performed experiments. J.R.W. synthesized luciferase substrates for bioluminescence imaging. Y.S., Y.W. and M.Z.L. analysed data. Y.S., Y.W. and M.Z.L. wrote the paper. M.M., T.A.K., and M.Z.L. provided supervision and obtained funding. All authors provided feedback on the manuscript.

## References

(1) Heffron, T. P. Small Molecule Kinase Inhibitors for the Treatment of Brain Cancer. J. Med. Chem. 2016, 59, 10030–10066. DOI: 10.1021/acs.jmedchem.6b00618

(2) Pearson, J. R. D.; Regad, T. Targeting cellular pathways in glioblastoma multiforme. Signal Transduct Target Ther 2017, 2, 17040. DOI: 10.1038/sigtrans.2017.40

(3) Smalley, K. S. M.; Forsyth, P. A. The Blood Brain Barrier and BRAF inhibitors: Implications for patients with melanoma brain metastases. Pharmacol. Res. 2018, 135, 265–267. DOI: 10.1016/j.phrs.2017.11.013

(4) Venur, V. A.; Ahluwalia, M. S. Targeted Therapy in Brain Metastases: Ready for Primetime Am Soc Clin Oncol Educ Book 2016, 35, e123–30. DOI: 10.1200/EDBK_100006

(5) Benn, C. L.; Dawson, L. A. Clinically Precedented Protein Kinases: Rationale for Their Use in Neurodegenerative Disease. Front Aging Neurosci 2020, 12, 242. DOI: 10.3389/fnagi.2020.00242

(6) Chico, L. K.; Van Eldik, L. J.; Watterson, D. M. Targeting protein kinases in central nervous system disorders. Nat Rev Drug Discov 2009, 8, 892–909. DOI: 10.1038/nrd2999

(7) Matsuda, S.; Ikeda, Y.; Murakami, M.; Nakagawa, Y.; Tsuji, A.; Kitagishi, Y. Roles of PI3K/AKT/GSK3 Pathway Involved in Psychiatric Illnesses. Diseases 2019, 7, E22. DOI: 10.3390/diseases7010022

(8) Zhang, D.; Hop, C. E. C. A.; Patilea-Vrana, G.; Gampa, G.; Seneviratne, H. K.; Unadkat, J. D.; Kenny, J. R.; Nagapudi, K.; Di, L.; Zhou, L.; Zak, M.; Wright, M. R.; Bumpus, N. N.; Zang, R.; Liu, X.; Lai, Y.; Khojasteh, S. C. Drug Concentration Asymmetry in Tissues and Plasma for Small Molecule-Related Therapeutic Modalities. Drug Metab. Dispos. 2019, 47, 1122–1135. DOI: 10.1124/dmd.119.086744

(9) Di, L.; Kerns, E. H.; Carter, G. T. Strategies to assess blood-brain barrier penetration. Expert Opin Drug Discov 2008, 3, 677–687. DOI: 10.1517/17460441.3.6.677

(10) Da Ros, M.; De Gregorio, V.; Iorio, A. L.; Giunti, L.; Guidi, M.; de Martino, M.; Genitori, L.; Sardi, I. Glioblastoma Chemoresistance: The Double Play by Microenvironment and Blood-Brain Barrier. Int J Mol Sci 2018, 19, 2879. DOI: 10.3390/ijms19102879;

(10a) Stocco, M. R.; Tyndale, R. F. Cytochrome P450 enzymes and metabolism of drugs and neurotoxins within the mammalian brain. Adv Pharmacol 2022, 95, 73–106. DOI: 10.1016/bs.apha.2022.04.003

(11) Onken, J.; Torka, R.; Korsing, S.; Radke, J.; Krementeskaia, I.; Nieminen, M.; Bai, X.; Ullrich, A.; Heppner, F.; Vajkoczy, P. Inhibiting receptor tyrosine kinase AXL with small molecule inhibitor BMS-777607 reduces glioblastoma growth, migration, and invasion in vitro and in vivo. Oncotarget 2016, 7, 9876–9889. DOI: 10.18632/oncotarget.7130

(12) Smith, D. A.; Rowland, M. Intracellular and Intraorgan Concentrations of Small Molecule Drugs: Theory, Uncertainties in Infectious Diseases and Oncology, and Promise. Drug Metab. Dispos. 2019, 47, 665–672. DOI: 10.1124/dmd.118.085951

(13) Greenwald, E. C.; Mehta, S.; Zhang, J. Genetically Encoded Fluorescent Biosensors Illuminate the Spatiotemporal Regulation of Signaling Networks. Chem Rev 2018, 118, 11707–11794. DOI: 10.1021/acs.chemrev.8b00333

(14) Liu, S.; Su, Y.; Lin, M. Z.; Ronald, J. A. Brightening up Biology: Advances in Luciferase Systems for in Vivo Imaging. ACS Chem Biol 2021, 16, 2707–2718. DOI: 10.1021/acschembio.1c00549;

(14a) Rice, B. W.; Cable, M. D.; Nelson, M. B. In vivo imaging of light-emitting probes. J Biomed Opt 2001, 6, 432–440. DOI: 10.1117/1.1413210

(15) Zhang, L.; Lee, K. C.; Bhojani, M. S.; Khan, A. P.; Shilman, A.; Holland, E. C.; Ross, B. D.; Rehemtulla, A. Molecular imaging of Akt kinase activity. Nat Med 2007, 13, 1114–1119. DOI: 10.1038/nm1608

(16) Zhang, L.; Virani, S.; Zhang, Y.; Bhojani, M. S.; Burgess, T. L.; Coxon, A.; Galban, C. J.; Ross, B. D.; Rehemtulla, A. Molecular imaging of c-Met tyrosine kinase activity. Anal. Biochem. 2011, 412, 1–8. DOI: 10.1016/j.ab.2011.01.028

(17) Su, Y.; Walker, J. R.; Park, Y.; Smith, T. P.; Liu, L. X.; Hall, M. P.; Labanieh, L.; Hurst, R.; Wang, D. C.; Encell, L. P.; Kim, N.; Zhang, F.; Kay, M. A.; Casey, K. M.; Majzner, R. G.; Cochran, J. R.; Mackall, C. L.; Kirkland, T. A.; Lin, M. Z. Novel NanoLuc substrates enable bright two-population bioluminescence imaging in animals. Nat Methods 2020, 17, 852–860. DOI: 10.1038/s41592-020-0889-6

(18) Chang, C.; Chan, A.; Lin, X.; Higuchi, T.; Terrovitis, J.; Afzal, J. M.; Rittenbach, A.; Sun, D.; Vakrou, S.; Woldemichael, K.; O’Rourke, B.; Wahl, R.; Pomper, M.; Tsui, B.; Abraham, M. R. Cellular bioenergetics is an important determinant of the molecular imaging signal derived from luciferase and the sodium-iodide symporter. Circ. Res. 2013, 112, 441–450. DOI: 10.1161/CIRCRESAHA.112.273375

(19) Hung, Y. P.; Teragawa, C.; Kosaisawe, N.; Gillies, T. E.; Pargett, M.; Minguet, M.; Distor, K.; Rocha-Gregg, B. L.; Coloff, J. L.; Keibler, M. A.; Stephanopoulos, G.; Yellen, G.; Brugge, J. S.; Albeck, J. G. Akt regulation of glycolysis mediates bioenergetic stability in epithelial cells. Elife 2017, 6, e27293. DOI: 10.7554/eLife.27293

(20) Papa, S.; Choy, P. M.; Bubici, C. The ERK and JNK pathways in the regulation of metabolic reprogramming. Oncogene 2019, 38, 2223–2240. DOI: 10.1038/s41388-018-0582-8

(21) Dranchak, P.; MacArthur, R.; Guha, R.; Zuercher, W. J.; Drewry, D. H.; Auld, D. S.; Inglese, J. Profile of the GSK published protein kinase inhibitor set across ATP-dependent and-independent luciferases: implications for reporter-gene assays. PLoS One 2013, 8, e57888. DOI: 10.1371/journal.pone.0057888

(22) Thorne, N.; Shen, M.; Lea, W. A.; Simeonov, A.; Lovell, S.; Auld, D. S.; Inglese, J. Firefly luciferase in chemical biology: a compendium of inhibitors, mechanistic evaluation of chemotypes, and suggested use as a reporter. Chem. Biol. 2012, 19, 1060–1072. DOI: 10.1016/j.chembiol.2012.07.015

(23) Ryan, M. B.; Corcoran, R. B. Therapeutic strategies to target RAS-mutant cancers. Nat Rev Clin Oncol 2018, 15, 709–720. DOI: 10.1038/s41571-018-0105-0

(24) Dixon, A. S.; Schwinn, M. K.; Hall, M. P.; Zimmerman, K.; Otto, P.; Lubben, T. H.; Butler, B. L.; Binkowski, B. F.; Machleidt, T.; Kirkland, T. A.; Wood, M. G.; Eggers, C. T.; Encell, L. P.; Wood, K. V. NanoLuc Complementation Reporter Optimized for Accurate Measurement of Protein Interactions in Cells. ACS Chem Biol 2016, 11, 400–408. DOI: 10.1021/acschembio.5b00753

(25) Pennell, S.; Westcott, S.; Ortiz-Lombardía, M.; Patel, D.; Li, J.; Nott, T. J.; Mohammed, D.; Buxton, R. S.; Yaffe, M. B.; Verma, C.; Smerdon, S. J. Structural and functional analysis of phosphothreonine-dependent FHA domain interactions. Structure 2010, 18, 1587–1595. DOI: 10.1016/j.str.2010.09.014

(26) Gonzalez, F. A.; Raden, D. L.; Davis, R. J. Identification of substrate recognition determinants for human ERK1 and ERK2 protein kinases. J. Biol. Chem. 1991, 266, 22159–22163.

(27) Lu, P. J.; Zhou, X. Z.; Shen, M.; Lu, K. P. Function of WW domains as phosphoserine-or phosphothreonine-binding modules. Science 1999, 283, 1325–1328. DOI: 10.1126/science.283.5406.1325

(28) Verdecia, M. A.; Bowman, M. E.; Lu, K. P.; Hunter, T.; Noel, J. P. Structural basis for phosphoserine-proline recognition by group IV WW domains. Nat Struct Biol 2000, 7, 639–643. DOI: 10.1038/77929

(29) Chu, J.; Oh, Y.; Sens, A.; Ataie, N.; Dana, H.; Macklin, J. J.; Laviv, T.; Welf, E. S.; Dean, K. M.; Zhang, F.; Kim, B. B.; Tang, C. T.; Hu, M.; Baird, M. A.; Davidson, M. W.; Kay, M. A.; Fiolka, R.; Yasuda, R.; Kim, D. S.; Ng, H. L.; Lin, M. Z. A bright cyan-excitable orange fluorescent protein facilitates dual-emission microscopy and enhances bioluminescence imaging in vivo. Nat Biotechnol 2016, 34, 760–767. DOI: 10.1038/nbt.3550

(30) Arvanitis, C. D.; Ferraro, G. B.; Jain, R. K. The blood-brain barrier and blood-tumour barrier in brain tumours and metastases. Nat Rev Cancer 2020, 20, 26–41. DOI: 10.1038/s41568-019-0205-x

(31) Alieva, M.; Leidgens, V.; Riemenschneider, M. J.; Klein, C. A.; Hau, P.; van Rheenen, J. Intravital imaging of glioma border morphology reveals distinctive cellular dynamics and contribution to tumor cell invasion. Sci Rep 2019, 9, 2054. DOI: 10.1038/s41598-019-38625-4

(32) de Gooijer, M. C.; Zhang, P.; Weijer, R.; Buil, L. C. M.; Beijnen, J. H.; van Tellingen, O. The impact of P-glycoprotein and breast cancer resistance protein on the brain pharmacokinetics and pharmacodynamics of a panel of MEK inhibitors. Int. J. Cancer 2018, 142, 381–391. DOI: 10.1002/ijc.31052

(33) Papale, A.; Morella, I. M.; Indrigo, M. T.; Bernardi, R. E.; Marrone, L.; Marchisella, F.; Brancale, A.; Spanagel, R.; Brambilla, R.; Fasano, S. Impairment of cocaine-mediated behaviours in mice by clinically relevant Ras-ERK inhibitors. Elife 2016, 5, e17111. DOI: 10.7554/eLife.17111

(34) Gilmartin, A. G.; Bleam, M. R.; Groy, A.; Moss, K. G.; Minthorn, E. A.; Kulkarni, S. G.; Rominger, C. M.; Erskine, S.; Fisher, K. E.; Yang, J.; Zappacosta, F.; Annan, R.; Sutton, D.; Laquerre, S. G. GSK1120212 (JTP-74057) is an inhibitor of MEK activity and activation with favorable pharmacokinetic properties for sustained in vivo pathway inhibition. Clin Cancer Res 2011, 17, 989–1000. DOI: 10.1158/1078-0432.CCR-10-2200

(35) Vaidhyanathan, S.; Mittapalli, R. K.; Sarkaria, J. N.; Elmquist, W. F. Factors influencing the CNS distribution of a novel MEK-1/2 inhibitor: implications for combination therapy for melanoma brain metastases. Drug Metab. Dispos. 2014, 42, 1292–1300. DOI: 10.1124/dmd.114.058339

(36) Yamaguchi, T.; Kakefuda, R.; Tajima, N.; Sowa, Y.; Sakai, T. Antitumor activities of JTP-74057 (GSK1120212), a novel MEK1/2 inhibitor, on colorectal cancer cell lines in vitro and in vivo. Int J Oncol 2011, 39, 23–31. DOI: 10.3892/ijo.2011.1015

(37) Allen, M.; Bjerke, M.; Edlund, H.; Nelander, S.; Westermark, B. Origin of the U87MG glioma cell line: Good news and bad news. Sci Transl Med 2016, 8, 354re3. DOI: 10.1126/scitranslmed.aaf6853

(38) Germann, U. A.; Furey, B. F.; Markland, W.; Hoover, R. R.; Aronov, A. M.; Roix, J. J.; Hale, M.; Boucher, D. M.; Sorrell, D. A.; Martinez-Botella, G.; Fitzgibbon, M.; Shapiro, P.; Wick, M. J.; Samadani, R.; Meshaw, K.; Groover, A.; DeCrescenzo, G.; Namchuk, M.; Emery, C. M.; Saha, S.; Welsch, D. J. Targeting the MAPK Signaling Pathway in Cancer: Promising Preclinical Activity with the Novel Selective ERK1/2 Inhibitor BVD-523 (Ulixertinib). Mol Cancer Ther 2017, 16, 2351–2363. DOI: 10.1158/1535-7163.MCT-17-0456;

(38a) Kessler, L.; Wu, T.; Guo, X.; Chen, J.; Hansen, R.; Thach, S. L. C.; Darjania, L.; Li, S. S.; Yu, K.; Ely, T. KO-947, a potent and selective ERK inhibitor with slow dissociation kinetics Poster ENA 2016,;

(b) Singh, M.; Nayyar, N.; Brastianos, P. DDRE-02. THERAPEUTIC TARGETING OF BRAIN METASTASIS WITH ERK INHIBITOR LY3214996 USING A NOVEL IN VIVO MODEL OF LUNG-TO-BRAIN METASTASIS Neuro-Oncology 2020, 22, ii61.

(39) Gupta, M.; Lee, H. J.; Barden, C. J.; Weaver, D. F. The Blood-Brain Barrier (BBB) Score. J. Med. Chem. 2019, 62, 9824–9836. DOI: 10.1021/acs.jmedchem.9b01220

(40) Jousma, E.; Rizvi, T. A.; Wu, J.; Janhofer, D.; Dombi, E.; Dunn, R. S.; Kim, M. O.; Masters, A. R.; Jones, D. R.; Cripe, T. P.; Ratner, N. Preclinical assessments of the MEK inhibitor PD-0325901 in a mouse model of Neurofibromatosis type 1. Pediatr Blood Cancer 2015, 62, 1709–1716. DOI: 10.1002/pbc.25546

(41) Nyati, S.; Young, G.; Ross, B. D.; Rehemtulla, A. Quantitative and Dynamic Imaging of ATM Kinase Activity by Bioluminescence Imaging. Methods Mol Biol 2017, 1599, 97–111. DOI: 10.1007/978-1-4939-6955-5_8;

(41a) Khan, A. P.; Contessa, J. N.; Nyati, M. K.; Ross, B. D.; Rehemtulla, A. Molecular imaging of epidermal growth factor receptor kinase activity. Anal. Biochem. 2011, 417, 57–64. DOI: 10.1016/j.ab.2011.05.040

(42) Leten, C.; Struys, T.; Dresselaers, T.; Himmelreich, U. In vivo and ex vivo assessment of the blood brain barrier integrity in different glioblastoma animal models. J Neurooncol 2014, 119, 297–306. DOI: 10.1007/s11060-014-1514-2

(43) Brighi, C.; Reid, L.; Genovesi, L. A.; Kojic, M.; Millar, A.; Bruce, Z.; White, A. L.; Day, B. W.; Rose, S.; Whittaker, A. K.; Puttick, S. Comparative study of preclinical mouse models of high-grade glioma for nanomedicine research: the importance of reproducing blood-brain barrier heterogeneity. Theranostics 2020, 10, 6361–6371. DOI: 10.7150/thno.46468

(44) Zhang, J. F.; Liu, B.; Hong, I.; Mo, A.; Roth, R. H.; Tenner, B.; Lin, W.; Zhang, J. Z.; Molina, R. S.; Drobizhev, M.; Hughes, T. E.; Tian, L.; Huganir, R. L.; Mehta, S.; Zhang, J. An ultrasensitive biosensor for high-resolution kinase activity imaging in awake mice. Nat Chem Biol 2021, 17, 39–46. DOI: 10.1038/s41589-020-00660-y

(45) Morciano, G.; Sarti, A. C.; Marchi, S.; Missiroli, S.; Falzoni, S.; Raffaghello, L.; Pistoia, V.; Giorgi, C.; Di Virgilio, F.; Pinton, P. Use of luciferase probes to measure ATP in living cells and animals. Nat Protoc 2017, 12, 1542–1562. DOI: 10.1038/nprot.2017.052

(46) Zhou, X. X.; Chung, H. K.; Lam, A. J.; Lin, M. Z. Optical control of protein activity by fluorescent protein domains. Science 2012, 338, 810–814. DOI: 10.1126/science.1226854

