## Supplementary Information for "Kinase-modulated bioluminescent indicators enable noninvasive imaging of drug activity in the brain"

### Table S1. Alanine scanning on SmBiT

| | WT (SmBiT) | V158A | T159A | G160A | **Y161A** | R162A | **L163A** | F164A | E165A | E166A | I167A | L168A |
|---|---|---|---|---|---|---|---|---|---|---|---|---|
| Fold change | 1.7 | 1.9 | 2.7 | 2.0 | **3.3** | 1.0 | **3.7** | 2.7 | 1.2 | 1.2 | 2.1 | 1.5 |
| Relative brightness (normalized to WT) | 1.0 | 0.61 | 0.43 | 0.038 | **0.35** | 0.0007 | **0.22** | 0.024 | 1.5 | 1.7 | 0.51 | 1.2 |

* Numbering based on NanoLuc sequence

### Table S2. Mutations on SmBiT sites 161 and 163

| | WT (SmBiT) | Y161A | Y161C | Y161L | L163A | L163K | **L163E** |
|---|---|---|---|---|---|---|---|
| Fold change | 1.7 | 3.3 | 3.4 | 3.3 | 3.7 | 2.4 | **3.7** |
| Relative brightness (normalized to WT) | 1.0 | 0.35 | 0.21 | 0.14 | 0.22 | 0.33 | **0.07** |

### Table S3. Screening of N-terminal fusions of fluorescent proteins

| Constructs | Fold change (+1 $\mu$M PD0325901) | Relative brightness(normalized to KiMBI0.1$_{ERK}$) | Red peak wavelength (nm) | BRET efficiency (peak around 600 nm/ peak at 450 nm) |
|---|---|---|---|---|
| KiMBI0.1$_{ERK}$ CyOFP1($\Delta$C7) L($\Delta$N2) WW S($\Delta$C3) Sub(V-3S) | 14.5 | 1.0 | - | - |
| mScarlet L($\Delta$N2) WW S($\Delta$C3) Sub(V-3S) | 2.4 | 6.2 | 587 | 0.31 |
| mCyRFP4 L($\Delta$N2) WW S($\Delta$C3) Sub(V-3S) | 2.9 | 5.2 | 597 | 0.27 |
| **tKiMBI$_{ERK}$** tdTomato($\Delta$C9) L($\Delta$N2) WW S($\Delta$C3) Sub(V-3S) | **11.3** | **2.0** | **579** | **0.36** |
| CyOFP1 CyOFP1($\Delta$C7) L($\Delta$N2) WW S($\Delta$C3) Sub(V-3S) | 12.2 | 4.8 | 583 | 0.18 |
| mScarlet CyOFP1($\Delta$C7) L($\Delta$N2) WW S($\Delta$C3) Sub(V-3S) | 9.7 | 2.0 | 593 | 0.14 |
| mScarlet mNeonGreen L($\Delta$N2) WW S($\Delta$C3) Sub(V-3S) | 12.7 | 0.48 | 594 | 0.34 |

### Table S4 Predicted BBB scores of select ERK inhibitors

| | KO-947 | Ulixertinib | Temuterkib |
|---|---|---|---|
| BBB score | 3.41 | 2.94 | 2.69 |
| PSA (Å$^2$) | 74 | 90 | 117 |
| HBD | 2 | 4 | 1 |
| cLogP | 2.4 | 4.1 | 1.4 |
| cLogD | 2.9 | - | 2.1 |
| MW | 355 | 432 | 454 |

PSA: polar surface area; HBD: hydrogen bond donors

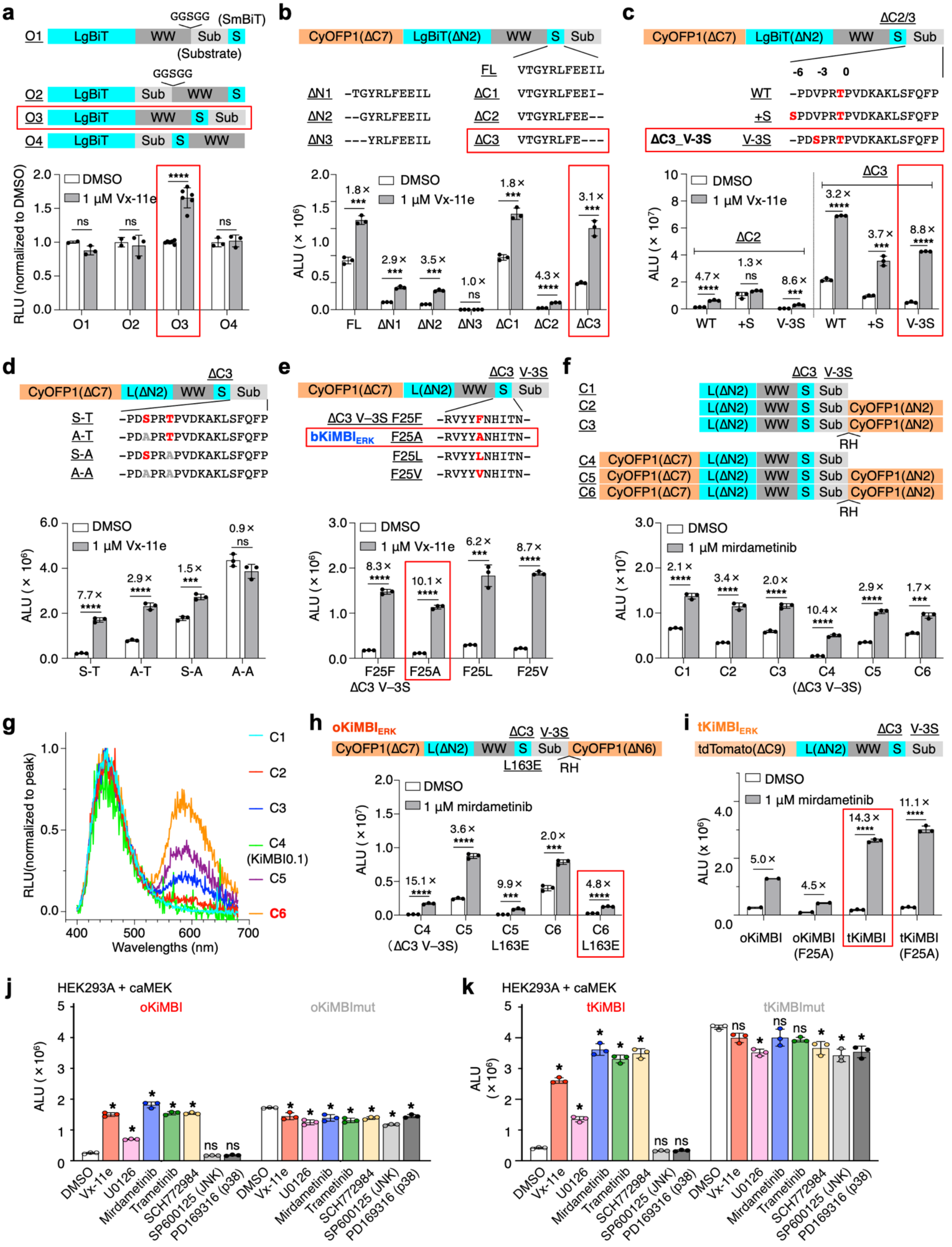
a
O1 LgBiT WW GGSGG Sub S (SmBiT) (Substrate)
O2 LgBiT Sub GGSGG WW S
O3 LgBiT WW S Sub
O4 LgBiT Sub S WW
RLU (normalized to DMSO)
DMSO
1 μM Vx-11e
b
CyOFP1(ΔC7) LgBiT(ΔN2) WW S Sub
FL VTGYRLFEEIL
ΔN1 -TGYRLFEEIL
ΔN2 --GYRLFEEIL
ΔN3 ---YRLFEEIL
ΔC1 VTGYRLFEEI-
ΔC2 VTGYRLFEE--
ΔC3 VTGYRLFE---
ALU (× 10⁶)
c
ΔC2/3
WT -PDVPRTPVDKAKLSFQFP
+S SPDVPRTPVDKAKLSFQFP
ΔC3_V-3S V-3S -PDSPRTPVDKAKLSFQFP
ALU (× 10⁷)
d
CyOFP1(ΔC7) L(ΔN2) WW S Sub
S-T -PDSPRTPVDKAKLSFQFP
A-T -PDAPRTPVDKAKLSFQFP
S-A -PDSPRAPVDKAKLSFQFP
A-A -PDAPRAPVDKAKLSFQFP
e
ΔC3 V–3S F25F -RVYYFNHITN-
bKiMBIERK F25A -RVYYANHITN-
F25L -RVYYLNHITN-
F25V -RVYYVNHITN-
f
C1 C2 C3 C4 C5 C6
RH
1 μM mirdametinib
(ΔC3 V–3S)
g
RLU(normalized to peak)
Wavelengths (nm)
C4 (KiMBI0.1)
h
oKiMBIERK
CyOFP1(ΔC7) L(ΔN2) WW S Sub CyOFP1(ΔN6)
L163E RH
i
tKiMBIERK
tdTomato(ΔC9) L(ΔN2) WW S Sub
oKiMBI oKiMBI (F25A) tKiMBI tKiMBI (F25A)
j
HEK293A + caMEK
oKiMBI oKiMBImut
DMSO Vx-11e U0126 Mirdametinib Trametinib SCH772984 SP600125 (JNK) PD169316 (p38)
k
HEK293A + caMEK
tKiMBI tKiMBImut

***Supplementary Fig. 1. Engineering, optimization, and characterization of KiMBIs. a,*** *Screen of LgBiT fused to WW, substrate peptide, and SmBiT (S) in various topolgies. HEK293A cells were transiently transfected with caMEK and each variant were treated with DMSO or MEK inhibitor and total bioluminescence assessed.* ***b****, Screening of SmBiT truncations to destabilize LgBiT-SmBiT reconstitution and increase inducibility.* ***c****, Introducing a second phosphorylation site to reduce background, yielding ⊿C3_V-3S. Potential phosphorylation sites for ERK kinase are labeled in red.* ***d****, Validation of phosphorylation sites in ⊿C3_V-3S. Serine/threonine to alanine mutations were shown in gray.* ***e****, Screening of mutations in WW domain for enhancing WW + pSub interactions, yielding bKiMBI$_{ERK}$.* ***f****, Introducing C-terminal CyOFP1 for red-shifting bioluminescence signal output.* ***g****, Bioluminescence spectra of KiMBI variants shown in* ***f****, and all spectra were normalized to the peak at around 450 nm.* ***h****, Introduction of mutations in SmBiT for improving fold induction of red-shifted KiMBI variants, yielding oKiMBI$_{ERK}$.* ***i****, Introduction of F25A mutation in WW for improving oKiMBI and tKiMBI. Error bars show s.d. Unpaired two-tailed Student's t-test was performed. ns, $p > 0.05$; *, $p < 0.05$; **, $p < 0.01$; ***, $p < 0.001$; **** $p < 0.0001$. Mean bioluminescence of HEK293A cells co-expressing caMEK and oKiMBI or oKiMBImut as negative control (****j****), or tKiMBI or tKiMBImut (****k****), in response to inhibitors Vx-11e (ERK, 1 µM), U0126 (MEK, 10 µM), mirdametinib (MEK, 1 µM), trametinib (MEK, 1 µM), SCH772984 (ERK, 1 µM), SP600125 (JNK,10 µM), and PD169316 (p38, 10 µM). Error bars show s.d. One-way ANOVA analysis ($p < 0.0001$) followed by Tukey's posthoc test was performed. ns, $p > 0.05$; *, $p < 0.05$.*

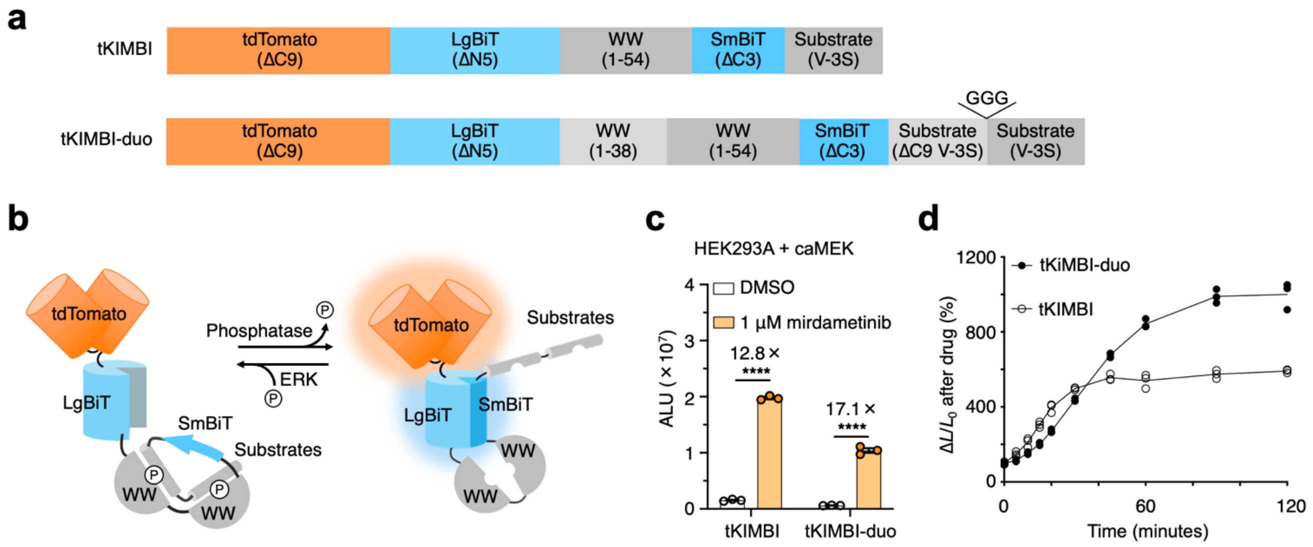


***Supplementary Fig. 2. Comparison between tKiMBI and tKIMBI-duo, a variant with repeated WW domains and substrates. a,*** *Domain structures of tKiMBI reporters.* ***b****, The proposed working mechanism of tKiMBI with two WW domains and substrates.* ***c****, Mean bioluminescence of tKiMBI variants in response to MEK inhibitor incubation in HEK293A cells transiently co-transfected with caMEK- and tKiMBI-encoded plasmids. Fold inductions of the signals between the DMSO- and inhibitor-treated samples were shown above the bar graphs. Error bars show s.d. Unpaired two-tailed Student's t-test was performed. ns, $p > 0.05$; **** $p < 0.0001$.* ***d****, Kinetics measurement of KiMBIs. Three technical replicates are shown for each group, and the result is representative of two biological replicates.*

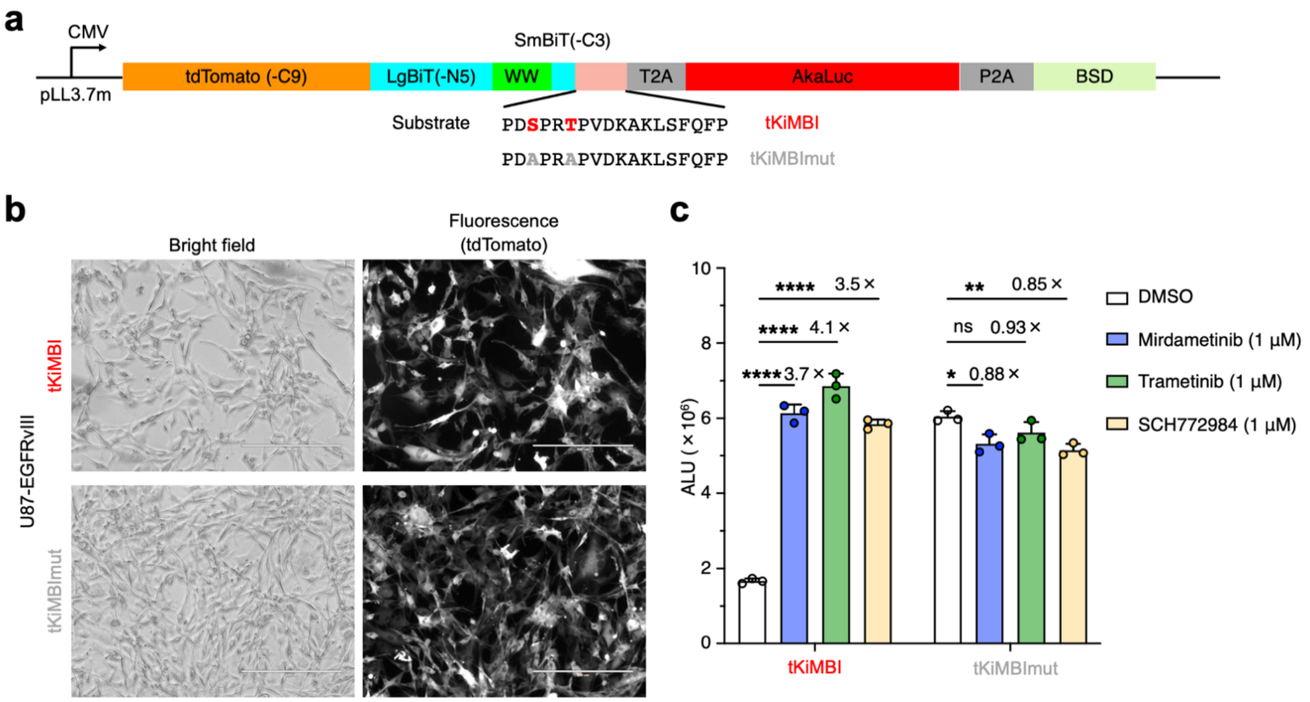


***Supplementary Fig. 3. In vitro characterization of U87-EGFRvIII stable reporter cell lines. a,*** *The domain structure of the tKiMBI (or tKiMBImut)-T2A-AkaLuc lentiviral plasmid for generating stable reporter cell lines. BSD, blasticidin-S deaminase, for antibiotic selection.* ***b****, Microscope images of U87-EGFRvIII reporter cell lines stably expressing tKiMBI or tKiMBImut.* ***c,*** *Mean bioluminescence of U87-EGFRvIII stable reporter cells in response to MEK-ERK inhibitors. Fold inductions of the signals between the DMSO- and inhibitor-treated samples were shown above the bar graphs. Error bars show s.d. Unpaired two-tailed Student's t-test. ns, $p > 0.05$; *, $p < 0.05$; **, $p < 0.01$; ***, $p < 0.001$; **** $p < 0.0001$.*

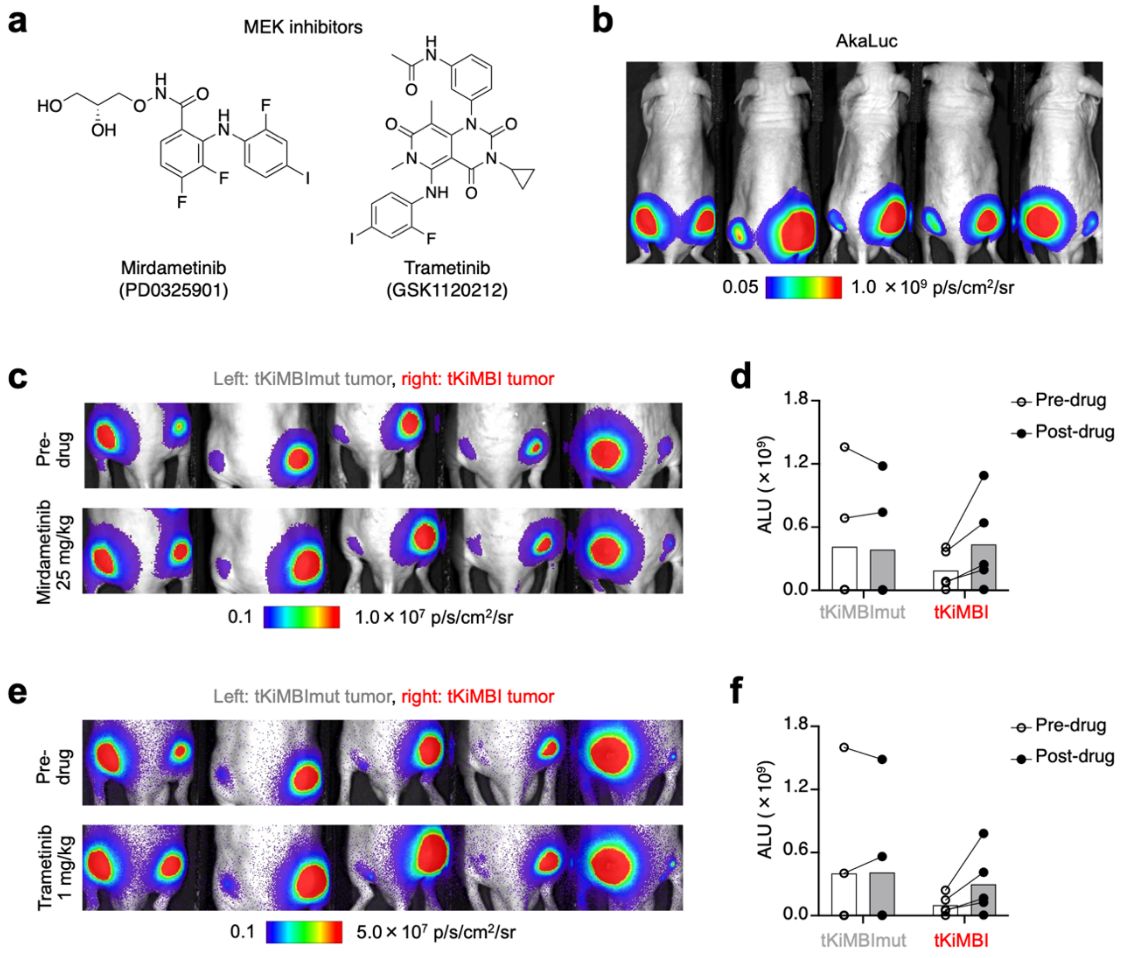


***Supplementary Fig. 4. Molecular imaging of ERK inhibition by MEK inhibitors in a subcutaneous tumor model. a,*** *Chemical structures of MEK inhibitors used in this experiment.* ***b****, Bioluminescence imaging of Akaluc at one week after implantation of U87-EGFRvIII reporter cells.* ***c-f,*** *(Raw data of* ***Fig. 2c,d****) U87-EGFRvIII tumor-bearing mice were sequentially treated (two days apart) with MEK inhibitors PD0325901 (****c-d****) and GSK1120212 (****e-f****) and were imaged with FFz injection 2 h after inhibitor injection.* ***c, e,*** *Bioluminescence imaging before and after inhibitor treatment.* ***d, f,*** *Total bioluminescence collected from the tumors. Each line represents a single tumor.*

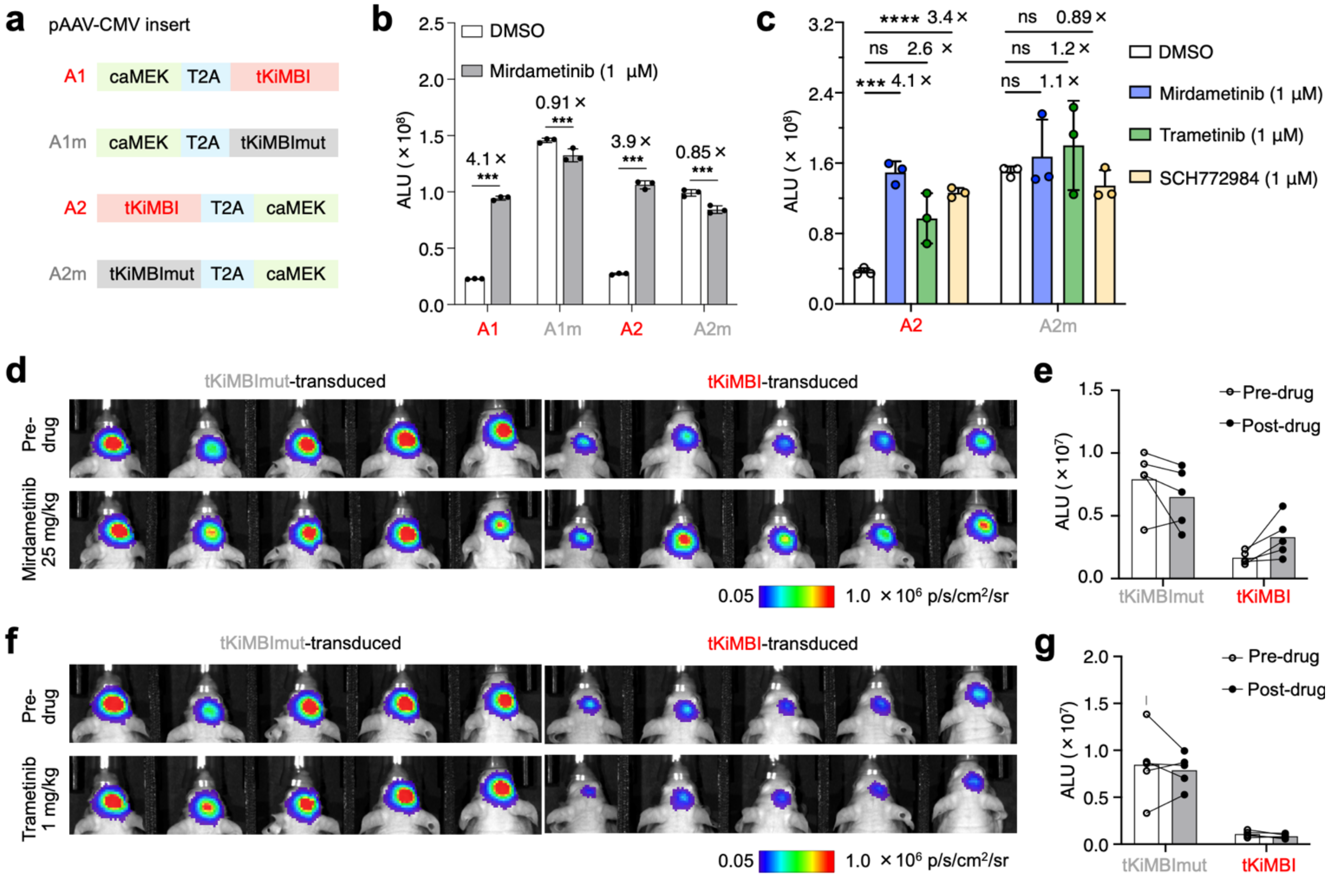


***Supplementary Fig. 5. In vitro characterization of AAV plasmids and bioluminescence imaging in AAV-infected mouse brain. a,*** *The domain structures of the tKiMBI- and caMEK- co-expressing AAV plasmids for in vivo gene delivery.* ***b, c****, Bioluminescence of HEK293A cells transiently transfected with AAV plasmids in response to MEK-ERK inhibitors incubation. Fold signal induction by each inhibitor relative to DMSO control is indicated above the bars. Error bars show s.d. Unpaired Student's t-test. ns, $p > 0.05$; ***, $p < 0.001$; **** $p < 0.0001$.* ***d-g****, (Raw data of* ***Fig. 3b,c)*** *Bioluminescence imaging of ERK inhibition in AAV infected mouse brain.* ***d, f,*** *Bioluminescence imaging before and after inhibitor treatment.* ***e, g,*** *Total bioluminescence collected from the brain tumors before and after inhibitor treatments. Each line represents an individual mouse.*

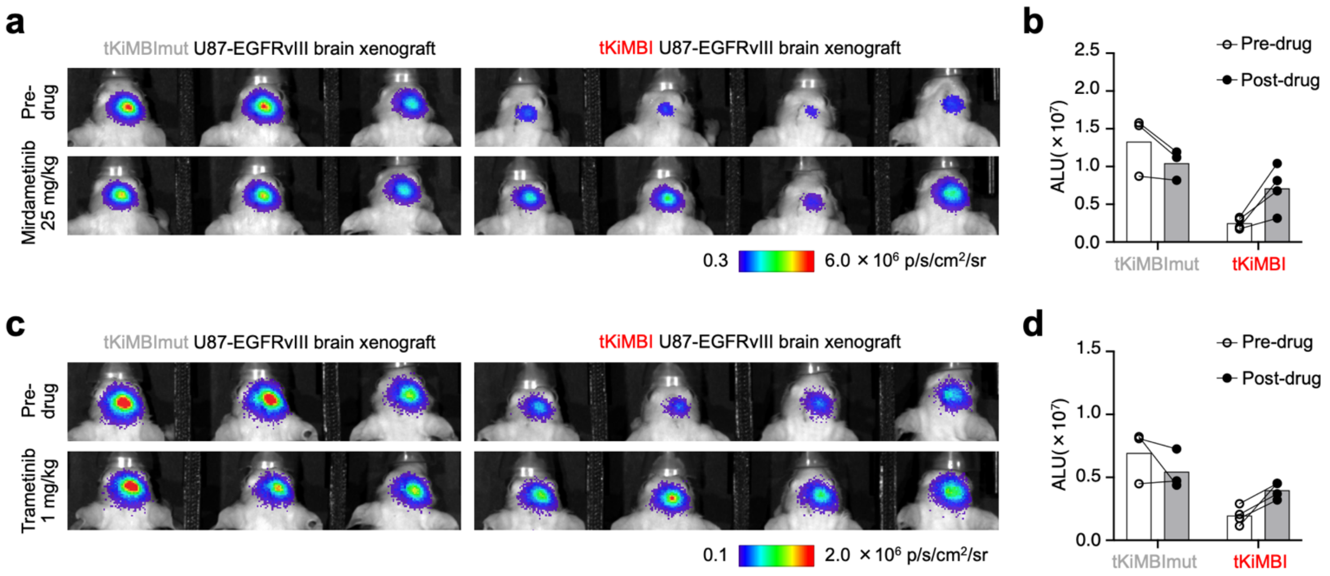


***Supplementary Fig. 6. Molecular imaging of ERK inhibition by MEK inhibitors in a brain tumor model. a-d, (****Raw data of* ***Fig. 3e****) of J:NU Mice with tKiMBI- or tKiMBImut-expressing U87-EGFRvIII tumor engrafted in the striatum were sequentially treated (two days apart) with PD0325901 (****a-b****) and GSK1120212 (****c-d****) and were imaged with CFz injection.* ***a*** *and* ***c,*** *Bioluminescence imaging before and after inhibitor treatment.* ***b*** *and* ***d,*** *Total bioluminescence collected from the brain tumors before and after inhibitor treatment.*

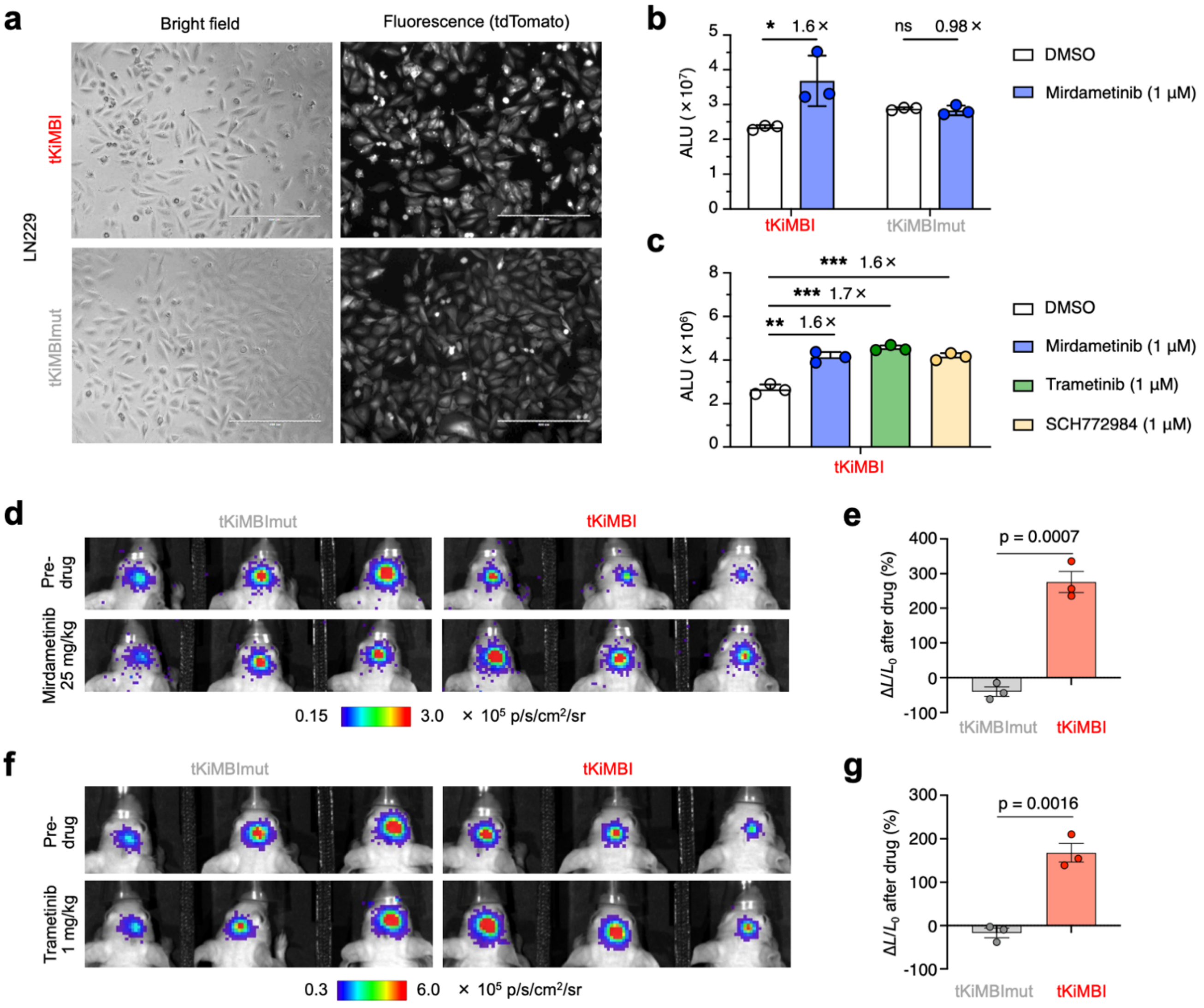


***Supplementary Fig. 7. Generation of a tKiMBI-expressing LN-229 cell line for molecular imaging of ERK inhibition in brain tumor xenografts. a,*** *Microscopic images of LN-229 reporter cell lines stably expressing tKiMBI or tKiMBImut.* ***b****, Mean bioluminescence of LN-229 stable reporter cell lines in response to PD0325901 incubation.* ***c****, Mean bioluminescence of tKiMBI-expressing LN-229 stable cell line in response to MEK-ERK inhibitors. Fold inductions of the signals between the DMSO- and inhibitor-treated samples were shown above the bar graphs. Error bars show s.d. Unpaired two-tailed Student's t-test. ns, $p > 0.05$; *, $p < 0.05$; **, $p < 0.01$; ***, $p < 0.001$.* ***d-g,*** *J:NU Mice with tKiMBI- or tKiMBImut-expressing LN-229 tumor engrafted in the striatum were sequentially treated (two days apart) with GSK1120212 (****d-e****) and PD0325901 (****f-g****) and were imaged with CFz injection.* ***d, f,*** *Bioluminescence imaging before and after inhibitor treatment.* ***e, g,*** *The percentage change of bioluminescence signals collected before and after inhibitor treatment. P values, unpaired Student's t-test.*

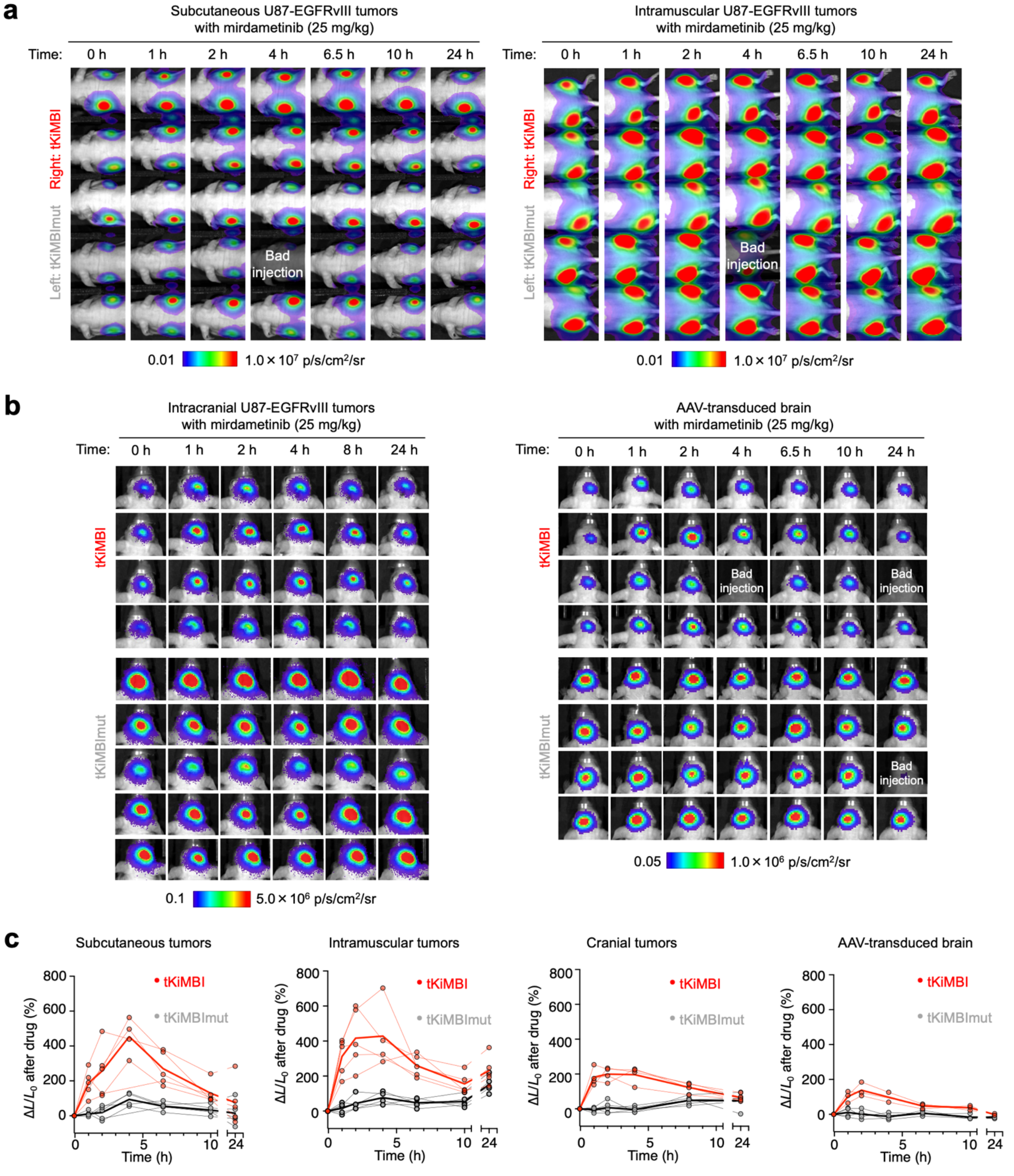


***Supplementary Fig. 8. Time-course molecular imaging of mirdametinib pharmacodynamics in target tissues. t****KiMBI and tKiMBImut responses over time after mirdametinib i.p. injection in J:NU mice bearing reporter-expressing U87-EGFRvIII extracranial tumors (**a**), bearing reporter-expressing U87-EGFRvIII cranial tumors (**b**, left), or transduced in the brain with reporter-expressing AAV (**a**, right)**.** These images constitute the entire dataset for **Fig. 4e**. Graphs of change in tKiMBI or tKiMBImut luminescence over time in the four tissues are shown in **c**. Thin lines, individual mice. Bold lines, mean change in signal over time for the entire group.*

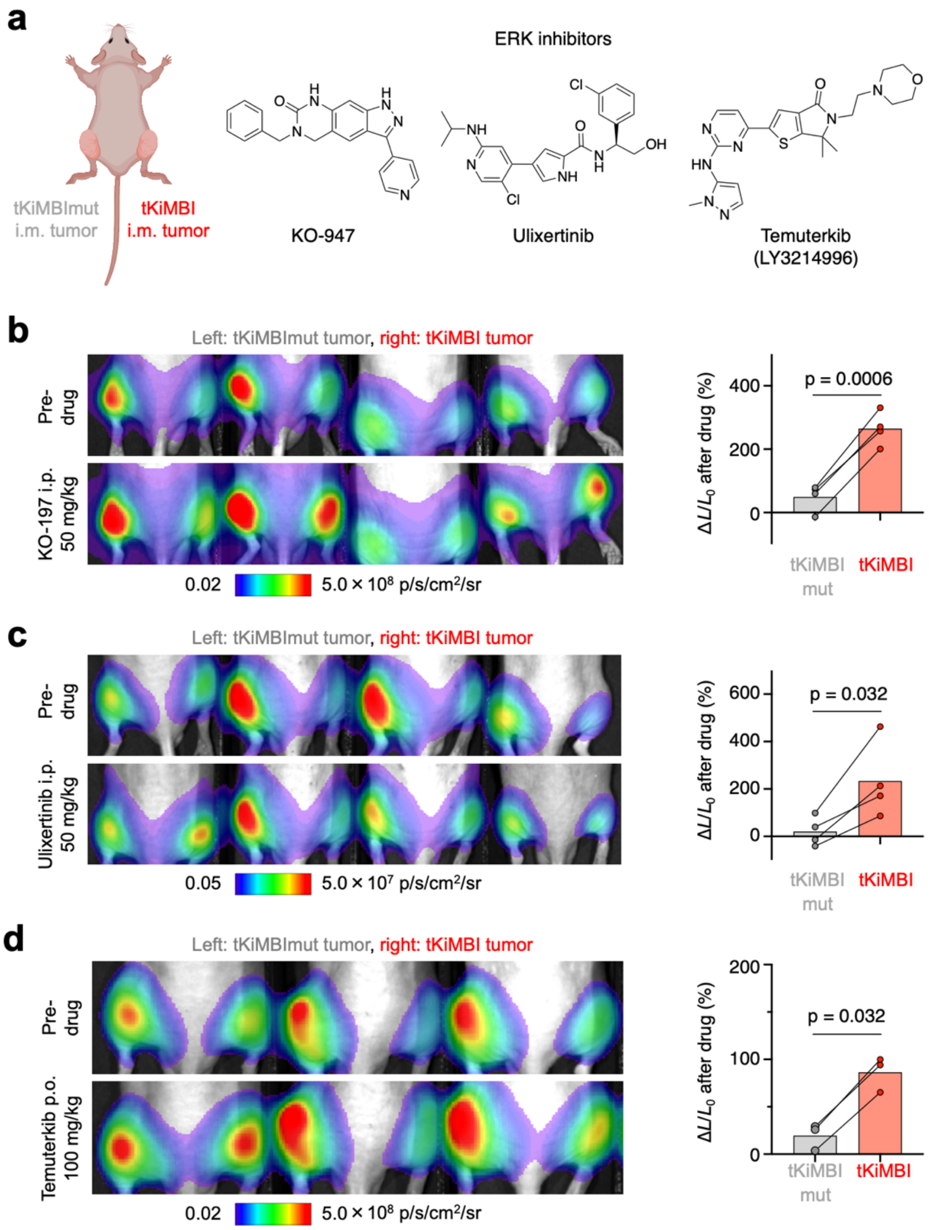


***Supplementary Fig. 9. Molecular imaging of ERK inhibitors in a tKiMBI-expressing intramuscular U87-EGFRvIII tumor cell xenograft model. a,*** *Scheme of intramuscular tumor xenograft model and chemical structures of ERK inhibitors used in this experiment.* ***c-d,*** *U87-EGFRvIII tumor-bearing mice were sequentially treated (two days apart) with ERK inhibitors KO-947 (**b**), ulixertinib (**c**) and LY3214996 (**d**) and were imaged with FFz injection 2 h after inhibitor injection. Left, bioluminescence imaging before and after inhibitor treatment. Right, the percentage change of bioluminescence signals collected before and after inhibitor treatment. Each line represents a single tumor. P values, paired two-tailed Student's t-test.*

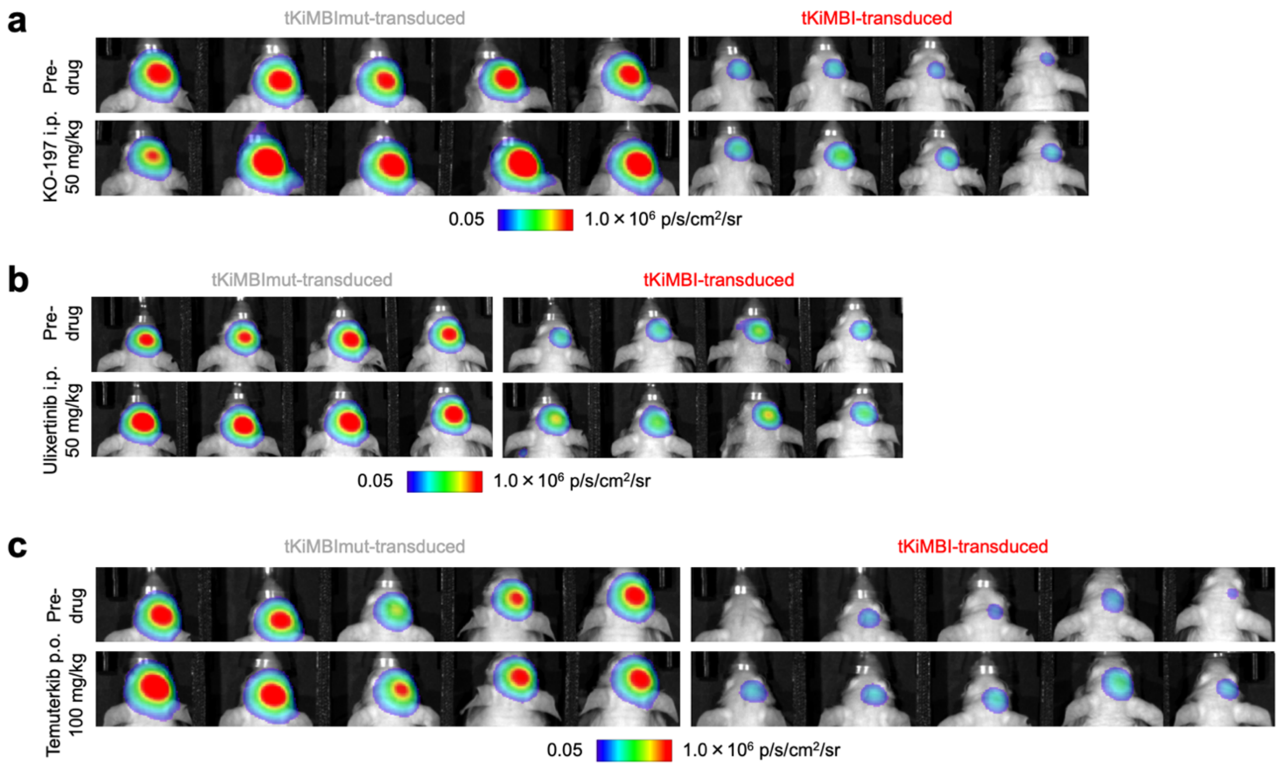


***Supplementary Fig. 10. Molecular imaging of ERK inhibitors in an AAV-infected mouse brain model. a-c,*** *(Raw images of Fig. 5b,c) J:NU Mice with tKiMBI- or tKiMBImut-expressing in the striatum after AAV transduction were sequentially treated (at least two days apart) with KO-947 (**a**), ulixertinib (**b**) and LY3214996 (**c**) and were imaged with CFz injection. Bioluminescence images were collected before and after inhibitor treatment.*
